# Antiproliferative Activity, Phytochemistry, Network Pharmacology, Molecular Docking and Gene Expression Analysis of *Maerua edulis* Extracts against Human Cervical Cancer Cell Line

**DOI:** 10.1101/2025.01.07.631450

**Authors:** Inyani John Lino Lagu, Rakita Letoluo, Sally Wambui Kamau, James Menni Kuria, Fredrick Mutie Musila, Vyacheslav Kungurtsev, Dorothy W. Nyamai, Sospeter Ngoci Njeru

## Abstract

*Maerua edulis* exhibits significant antiproliferative activity against HeLa cells, with hexane and ethyl acetate extracts showing IC_50_ values of 0.02% and 47.42 ***µ***g/mL, respectively. Gas Chromatography-Mass Spectrometry (GC-MS) analysis identified key phytochemicals such as diisooctyl phthalate, squalene, and stigmasta-3,5-diene, which were associated with the regulation of apoptotic and cell cycle-related genes (BCL2, CDK2, TP53). Gene expression assays confirmed the modulation of these targets, suggesting the therapeutic potential of *M. edulis* in cervical cancer. Further in vivo studies are recommended to validate these findings and establish its safety profile.

## 1 Introduction

Cervical cancer is the fourth most common cancer among women in the world [62], and a leading cause of cancer deaths. Despite being highly preventable, 604,127 women were diagnosed with, and 341,831 women died from this cancer, globally in 2020 [72]. The global burden of cervical cancer is expected to increase to about 28.4 million cases by the year 2040, an increase of 47% compared to the cases reported in 2020 [75]. The highest burden of cervical cancer is reported in low- and middle-income countries (LMIC), with sub-Saharan Africa and Southeast Asia having the highest mortality rates compared to the rest of the world [60].

Cervical cancer can be treated with surgery, radiotherapy, chemotherapy, and immunotherapy [62, 43]. Although these treatments have exhibited efficacy, these strategies are often unsatisfactory and are characterized by adverse side effects [30]. These effects are attributed to the non-selective nature of chemotherapeutic drugs, which often target non-cancerous cells [49]. Moreover, metastatic cervical cancers pose a risk of relapse after remission following radio-chemotherapy treatment [59]. Another emerging obstacle is chemotherapy resistance, resulting in unpredictable resistance and unsatisfactory outcomes from anticancer and cytotoxic agents [62, 9].

Due to the complexity associated with conventional treatments, their limitations, and high costs, there is a need to evaluate and explore safe, efficacious, and affordable chemotherapeutic drugs [85, 11]. Phytotherapy is one such alternative that offers advantages over traditional therapies [18]. Medicinal plants and herbs have a long history of treating various diseases, including cancer, and therefore have become one of the main modalities in complementary and alternative medicine [11]. Various chemotherapeutic drugs derived from medicinal plants and used in their natural form or with structural modifications, such as paclitaxel, triterpenoid botulin, piperine, resveratrol, epicatechin, epigallocatechin gallate, and flavonoids such as quercetin, silibinin, and oncamex, have shown the ability to induce apoptosis in cancer cells [49, 18, 59].

Some ethnobotanical and scientific studies have identified and screened several medicinal plants and herbs with potential anticancer activities [11, 85, 9]. One such medicinal plant is *Maerua edulis* [60, 49]. *Maerua edulis* (Gilg & Gilg-Ben.) DeWolf, commonly referred to as the blue-leaved bush berry, is a small perennial shrub in the family *Capparaceae* [49]. It is an evergreen shrub, about three meters in height, with alternate, simple, fleshy, hairless leaves, with thick, swollen, tuberous roots [85].

*M. edulis* is indigenous to Kenya, South Africa, the Democratic Republic of Congo, Malawi, Uganda, Mozambique, Tanzania, Zambia, and Zimbabwe [49, 11]. Traditionally, this plant has been used to treat and manage various ailments, including eye infections, cough, tuberculosis, pain, allergies, infertility in women, rheumatic swelling, and sexually transmitted diseases [49, 11]. Sithole and Mukanganyama (2018) evaluated the antiproliferative activity of *M. edulis* on Jurkat-T cells. However, no scientific research has explored the antiproliferative and cytotoxic effects of this medicinal plant against human cervical disease model.

Network pharmacology (NP) is a new discipline that integrates computer science and biological s [60]. NP helps to identify new drug candidates and repurpose existing ones from intricate network models [49]. The study of medicinal plants poses unique challenges due to their complexity in composition, multiple targets, and therapeutic signaling pathways [11]. However, NP can elucidate the effects of medicinal herb formulations on complex diseases such as cancer, thereby predicting putative therapeutic phytochemicals, their targets and decipher modes of action. In this study, the active phytochemicals, targets, and related signaling pathways of the ethyl acetate and hexane fractions of *M. edulis* in the treatment of cervical cancer were explored. Key compounds and targets of *M. edulis* fractions acting on cervical cancer were obtained through GC-MS and network topology analysis, annotated by gene ontology and pathway enrichment, evaluated via molecular docking (MD), and confirmed through gene expression and functional assays analysis to provide a scientific basis for the clinical application of *M. edulis*.

## 2 Materials and Methods

### 2.1 Plant collection and preparation

Roots of *Maerua edulis* were collected from Embu County, Mbeere South sub-County, Mavuria ward (Latitude 0*^◦^*46‘27.0”S, Longitude 37*^◦^*40‘54.9”E). Plant identification and authentication were performed by a botanist at the University of Nairobi, and a voucher specimen number (NSN18) was given and deposited at the Kenyan National Museums - Eastern Africa Herbarium. The plant materials were air-dried at 25−30 *^◦^*C for three weeks, pulverized into fine powder, and stored in an airtight plastic bag at room temperature until further analysis.

### 2.2 Plant extraction and fractionation

The powdered plant material was subjected to cold solvent extraction using dichloromethane:methanol (DCM:MeOH, 1:1 ratio), following the protocol described by Lagu et al. (2024) [40]. Five hundred grams of powdered plant material were macerated in 1 liter of DCM:MeOH for 48 hours, filtered through Whatman No. 1 filter paper, and the process was repeated thrice to ensure exhaustive extraction of phytochemicals. The filtrates were pooled and concentrated using a rotary evaporator. The obtained crude extract was fractionated sequentially by solvent partitioning with solvents of increasing polarity (n-hexane, ethyl acetate, and distilled water) as described in prior studies [40, 56, 14, 31, 57]. The organic crude extract and solvent fractions were concentrated, reconstituted in 100% dimethyl sulfoxide (DMSO), and diluted to ensure a final DMSO concentration of 0.2% in the test sample. The aqueous fraction was reconstituted in physiological saline. All stock solutions were stored at −20 *^◦^*C until analysis.

### 2.3 Cell culture

#### 2.3.1 Cell lines and culture conditions

Human cervical cancer HeLa-229 cells and non-cancerous kidney epithelial cells (Vero-CCL) were obtained from the American Type Culture Collection (ATCC). The cells were cultured in Essential Modified Eagle Medium (EMEM) supplemented with 1% L-glutamine (200 mM), 10% fetal bovine serum (FBS), 1.5% sodium bicarbonate, 1% HEPES buffer (1 M), 1% penicillin-streptomycin, and 0.25 *µ*g*/*mL amphotericin B. Cells were maintained at 37 *^◦^*C in a humidified atmosphere with 5% CO_2_ to achieve 80% confluence.

#### 2.3.2 In-vitro antiproliferation and cytotoxicity testing

The antiproliferative activity of the total extract, ethyl acetate and water extract fractions (at a fixed concentration of 200 *µ*g*/*mL and hexane fraction (at 0.2%) were screened against HeLa-229 cells using the MTT assay [40, 31, 54]. HeLa cells were seeded at 1 × 10^4^ cells/well in a 96-well plate and grown overnight for attachment. Cells were then treated with the total extract and fractions for 48 hours. Doxorubicin (200 *µ*g*/*mL) and 0.2% DMSO were used as positive and negative controls, respectively. After incubation, 10 *µ*L of 5 mg/mL MTT solution was added to each well and incubated at 37 *^◦^*C for 4 hours. Subsequently, 100 *µ*L of 100% DMSO was added to dissolve the formazan crystals. Absorbance was measured at 570 nm with a reference wavelength of 720 nm, using a plate reader (Infinite M1000, Tecan). Treatments showing growth inhibition greater than 50% were considered active. Cell viability (%) was calculated as follows:

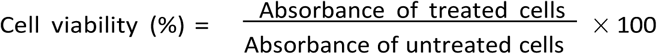

Ethyl acetate and hexane fractions of *M. edulis* exhibited robust growth inhibition of HeLa cells and were selected for further investigation. HeLa cells were treated with ethyl acetate fractions at concentrations of 2, 4, 8, 16, 32, and 64 *µ*g*/*mL and hexane fractions at 0.003, 0.006, 0.01, 0.025, 0.05, and 0.1%. The non-cancerous Vero cells were treated with the same fractions, at concentrations ranging from 32–1024 *µ*g*/*mL (ethyl acetate) and 0.008–0.25% (hexane), to assess safety and selectivity. The halfmaximal inhibitory concentration (IC_50_) and half-maximal cytotoxic concentration (CC_50_) values were calculated using non-linear regression in GraphPad Prism 8.4. Selectivity Index (SI) was determined using the formula:

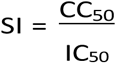

#### 2.3.3 Morphological study

HeLa cells were seeded at 1 × 10^4^ cells/well in a 96-well plate and treated with varying concentrations of ethyl acetate fractions (2–64 *µ*g*/*mL) and hexane fractions (0.003–0.1%) of *M. edulis* for 48 hours. Morphological changes were observed using an EVOSTM XL Core Imaging system (Thermo Scientific, USA) at 20x magnification [83].

#### 2.3.4 Wound healing assay

HeLa cells (1×10^5^ cells/well) were seeded in a 24-well plate and incubated for 24 hours to form a monolayer. A vertical wound was created using a sterile 200 *µ*L pipette tip. Cells were treated with ethyl acetate (47.42 *µ*g*/*mL) and hexane fractions (0.02%) of *M. edulis* and incubated for 48 hours. Images of the wounds were captured at 0, 24, and 48 hours using the EVOSTM XL Core Imaging system. The relative migration distance (RMD) was calculated as follows:

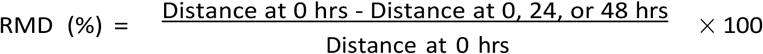

where the distance is computed within scratch.

### 2.4 Phytochemical analysis

#### 2.4.1 Qualitative phytochemical screening

Preliminary qualitative screening of phytochemical constituents was conducted to determine the presence or absence of alkaloids, glycosides, phenols, flavonoids, terpenoids, steroids, quinones, saponins, and tannins by visual inspection of color changes or precipitation in *M. edulis* ethyl acetate (EAF) and hexane (HF) fractions, following standard methods [53, 55, 5, 44].

#### 2.4.2 Semi-quantitative gas chromatography-mass spectrometry (GC-MS) analysis

The detection of bioactive compounds in *M. edulis* EAF and HF fractions was performed using a gas chromatography-mass spectrometry (GC-MS) system (Shimadzu GC-MS QP-2010SE) equipped with a low-polarity BPX5 capillary column (30 m × 0.25 mm × 0.25 µm film thickness). The oven temperature was programmed to 55°C for 1 min, increased by 10 °C/min to 280 °C, and held for 15 min. The injector temperature was set at 200 °C. Helium (99.999%) was used as the carrier gas at a flow rate of 1.08 mL/min. A sample volume of 1 µL (1% v/v) was injected in split mode (10:1 ratio). The ion source and interface temperatures were 200 °C and 250 °C, respectively. Mass spectra were collected at 70 eV in full scan mode (*m/z* 35–550). The GC-MS spectra were compared with entries in the NIST library database to identify the phytochemicals [53, 31].

### 2.5 In-silico analysis

#### 2.5.1 Drug-like screening and ADME-Tox profile

The PubChem canonical SMILES of GC-MS identified compounds in *M. edulis* EAF and HF were retrieved from the PubChem database ^1^. These were input into SwissADME^2^ and pkCSM^3^ to predict drug-likeness and ADME-Tox (absorption, distribution, metabolism, excretion, and toxicity) profiles. Drug-likeness predictions followed Lipinski’s Rule of Five, with thresholds including human intestinal absorption (HIA) ≥90%, blood-brain barrier (BBB) permeability, and central nervous system (CNS) activity [87, 12].

#### 2.5.2 Prediction of phytocompound-related gene targets

Potential target genes for *M. edulis* EAF and HF compounds were predicted using Swiss TargetPrediction^4^, Binding Database ^5^, and the Similarity Ensemble Approach (SEA) ^6^. A normalized fit score threshold of ≥0.1 for Swiss TargetPrediction and ≥0.9 for Binding Database was used [20]. Targets names were standardized using the UniProt database ^7^.

#### 2.5.3 Prediction of human cervical cancer-related target genes

Human cervical cancer-related genes were identified from the GeneCards ^8^, DisGeNET ^9^, OMIM ^10^ and Pharos ^11^ databases using a keyword search for “Cervical cancer” and thus yielded potential targets. Duplicates were removed to have a single disease target list. Overlapping targets between the disease- and phytocompound-targets were visualized using the Venn online tool ^12^ [90].

#### 2.5.4 Protein-protein interaction (PPI) network analysis

Protein-protein interactions were assessed using the STRING database ^13^, limiting the species to *Homo sapiens*. A confidence score of *>* 0.6 was set as the minimum interaction threshold. The PPI network was visualized and analyzed using Cytoscape v3.10 software. Hub genes were selected using CytoHubba’s degree algorithm with thresholds of Degree of ≥50 and Betweenness Centrality (BC) of ≥50 [48].

#### 2.5.5 Functional enrichment analysis

Gene Ontology (GO) and Kyoto Encyclopedia of Genes and Genomes (KEGG) enrichment analyses were performed using ShinyGO v0.77 ^14^. “Human” was selected as the species, with a false discovery rate (FDR) of ≤50 and *p <* 0.05. The top 15 GO terms (encompassing Biological Process, Molecular Function, Cellular Component) and KEGG pathways were visualized as histograms [46].

#### 2.5.6 Molecular docking

Protein 3D structures (in PDB format) were downloaded from the Protein Data Bank ^15^ for selected targets (e.g., CDK2: 4GCJ, TP53: 7LIN). The structures of *M. edulis* phytocompounds were retrieved from PubChem and converted to PDBQT format using Open Babel GUI v3.1.1. Molecular docking was performed using PyRx AutoDock VINA. Binding affinity thresholds were set at ≤ −7.5 kcal/mol for ethyl acetate fractions and ≤ −6.0 kcal/mol for hexane fractions. The results were visualized with Discovery Studio (version 2021 Client) [33].

### 2.6 Validation of gene targets by quantitative real-time PCR (RT-qPCR)

Gene expression analysis was conducted to validate molecular docking results. RNA was extracted from HeLa cells treated with *M. edulis* fractions at IC_50_ concentrations using the Total RNA Extraction Kit (Solarbio, Beijing, China). RNA concentration, quality and purity were checked using a NanoDrop spectrophotometer (Thermo Fisher). cDNA synthesis was performed with the SensiFAST cDNA synthesis kit (Brisbane, Australia). Gene expression levels for BCL2, CDK2, TP53, CASP9, and other targets were quantified using SensiFAST SYBR Lo-ROX Kit (Bioline, Australia) under the following conditions: 95*^◦^*C for 2 min, 40 cycles of 95 *^◦^*C for 5 s, 62 *^◦^*C for 10 s, and 72*^◦^*C for 20 s. Relative expression was calculated using the 2*^−^*^ΔCt^ method, with GAPDH as the reference gene.

### 2.7 Statistical analysis

Data are expressed as mean ± standard deviation (M ± SD). Statistical analyses were performed using GraphPad Prism v8.4 (San Diego, CA, USA). Comparisons were conducted via one-way ANOVA, followed by Tukey’s multiple comparison test. Results statistical significance was expressed as:

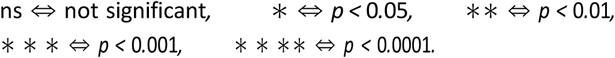

### 2.8 Ethical approval

This research was approved by the Kenya Medical Research Institute’s IRB and the Scientific and Ethics Review Unit (SERU) under approval number: KEMRI/SERU/CTMDR/104/4466.

## 3 Results

### 3.1 Antiproliferative activity of *M. edulis* extracts on HeLa cells

An initial antiproliferative screening of *M. edulis* extracts (crude, hexane, ethyl acetate, and water extracts) was undertaken on HeLa cells (*in vitro* model for cervical cancer) using the MTT assay at a fixed concentration of 200 *µ*g/mL (for crude-, ethyl acetate- and water-extracts) and 0.2% (V/V) for hexane extract. Ethyl acetate and hexane extracts exhibited significant (*p <* 0.0001) growth inhibition of HeLa cells relative to the negative control (NC; 0.2% DMSO), suppressing growth by more than 50% (our screening threshold) (Figure 3.1). *M. edulis* ethyl acetate and hexane extracts passed our cut-off and were therefore selected for concentration-dependent antiproliferative and cytotoxicity testing to determine their *IC*_50_ and *CC*_50_ values.

### 3.2 *M. edulis* ethyl acetate and hexane extracts exhibit concentration-dependent and selective bioactivity

Concentration-response studies of ethyl acetate and hexane extracts of *M. edulis* were performed to determine the concentrations that inhibit the growth of half of the treated HeLa cells (*IC*_50_) and that are cytotoxic to half of the treated non-cancerous Vero cells (*CC*_50_). HeLa and Vero cells were treated with ethyl acetate and hexane extracts at varying concentrations for 48 h (Figures 2 and 3). Ethyl acetate extract had *IC*_50_ and *CC*_50_ values of 47.42 ± 3.11 *µ*g*/*mL and 545.16 ± 6.76 *µ*g*/*mL against HeLa and Vero cells, respectively. Hexane extract exhibited *IC*_50_ and *CC*_50_ values of 0.02 ± 0.003% and 0.12 ± 0.01% against HeLa and Vero cells, respectively. Doxorubicin exhibited an *IC*_50_ value of 2.09 ± 1.35 *µ*g*/*mL and a *CC*_50_ value of 3.44 ± 1.00 *µ*g*/*mL. Therefore, the antiproliferative activity of tested extract fractions on HeLa cells and cytotoxicity effects on Vero cells were concentration-dependent.

**Fig. 1.**
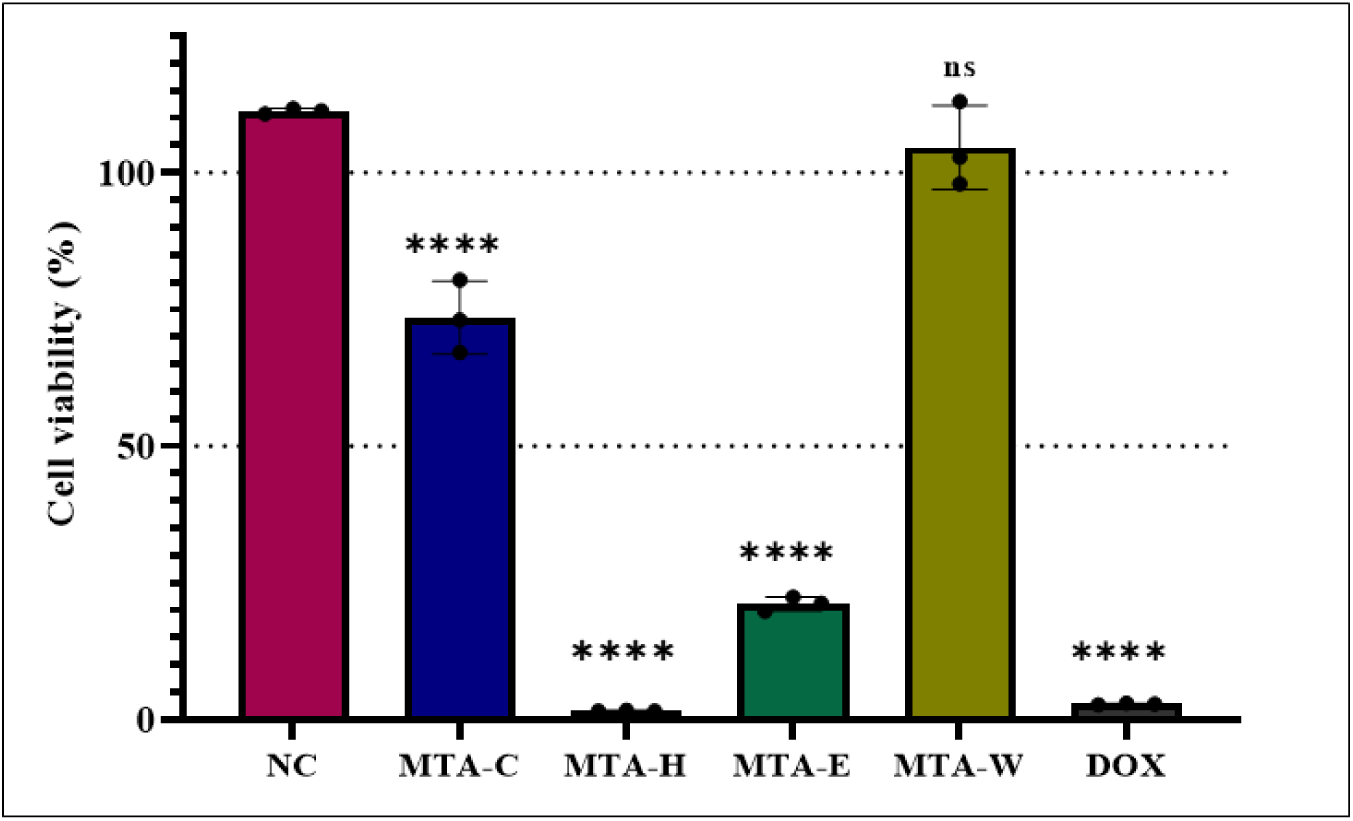
Antiproliferative activity results of *M. edulis* extracts on HeLa cells at a fixed concentration of 200 *µ*g/mL (total (DCM:MeOH) crude extract, ethyl acetate and water fractions) and 0.2% (V/V) for hexane fraction. Doxorubicin (200 *µ*g/mL) was used as a positive control, and 0.2% DMSO was used as a negative control (NC). Data are presented as Mean *±* S.D. of three independent experiments. **** represent, p *<*0.0001; ns, not significance. MTA-C, MTA-H, MTA-E, MTA-W, and Dox represent crude extract, hexane fraction, ethyl acetate fraction, water fractions of *M. edulis*, and doxorubicin, respectively.

**Fig. 2.**
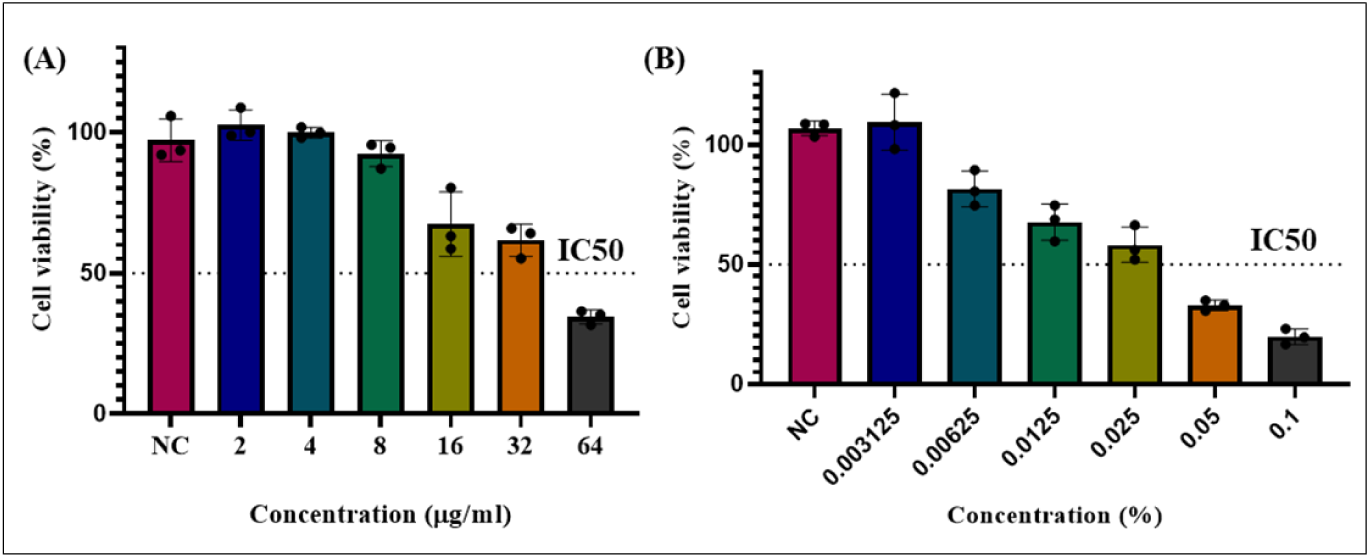
*In vitro* antiproliferative effects of *M. edulis* fractions at different concentrations against HeLa cells after 48 h of incubation with; (A) Ethyl acetate fraction and, (B) Hexane fractions of *M. edulis*. Data are represented as Mean ± S.D. of three independent experiments performed in triplicates. NC represents the negative control.

**Fig. 3.**
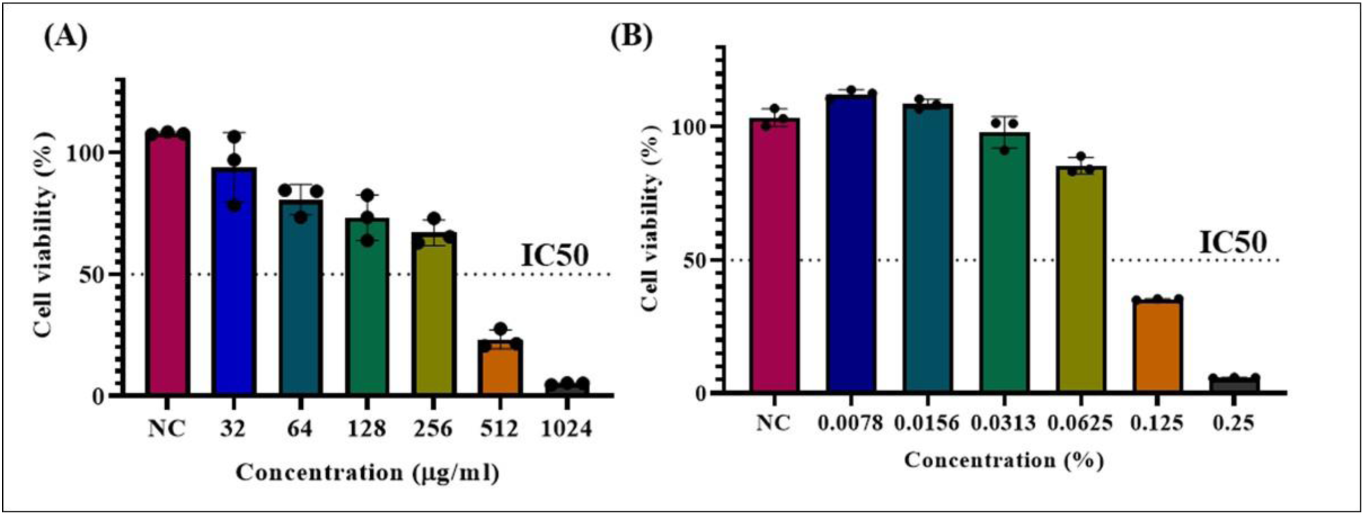
*In vitro* cytotoxicity effects of *M. edulis* fractions at different concentrations against Vero cells after 48 h of incubation with; (A) Ethyl acetate fraction and, (B) Hexane fractions of *M. edulis*. Data are represented as Mean ± S.D. of three independent experiments performed in triplicates. NC represents the negative control.

### 3.3 Selective index (SI) of *M. edulis* and doxorubicin

The selectivity index (SI) was calculated to evaluate the selectivity of *M. edulis* ethyl acetate and hexane fractions against cancer cells compared to non-cancerous cells. The SI values for ethyl acetate and hexane fractions were 11.5 and 6.32, respectively (Table **??**). These results imply that *M. edulis* ethyl acetate and hexane fractions have higher selectivity for cancerous cells with minimal cytotoxic effects on non-cancerous cells, indicating excellent safety of the plant extract fractions at the tested bioactive dose.

### 3.4 *M. edulis* ethyl acetate and hexane fractions induce morphological changes in HeLa cells

The antiproliferative effects of *M. edulis* ethyl acetate and hexane fractions were assessed by exposing HeLa cells to different concentrations of the extract fractions. Changes were observed under a microscope after 48 hours of exposure (Figures 4 and 5). In the negative control, HeLa cells displayed an angular shape and normal growth pattern, forming a patchy monolayer. In contrast, HeLa cells treated with both fractions of *M. edulis* showed significant morphological alterations, including reduced cell viability, loss of adhesion, a spherical shape, sporadic distribution, and dead cell debris. At the highest concentrations of ethyl acetate (64 *µ*g*/*mL) and hexane fractions (0.25%), complete disruption of cell morphology was observed. HeLa cells treated with doxorubicin (2.09 *µ*g*/*mL) (positive control) exhibited reduced cell viability and atypical morphological changes.

**Table 1:**
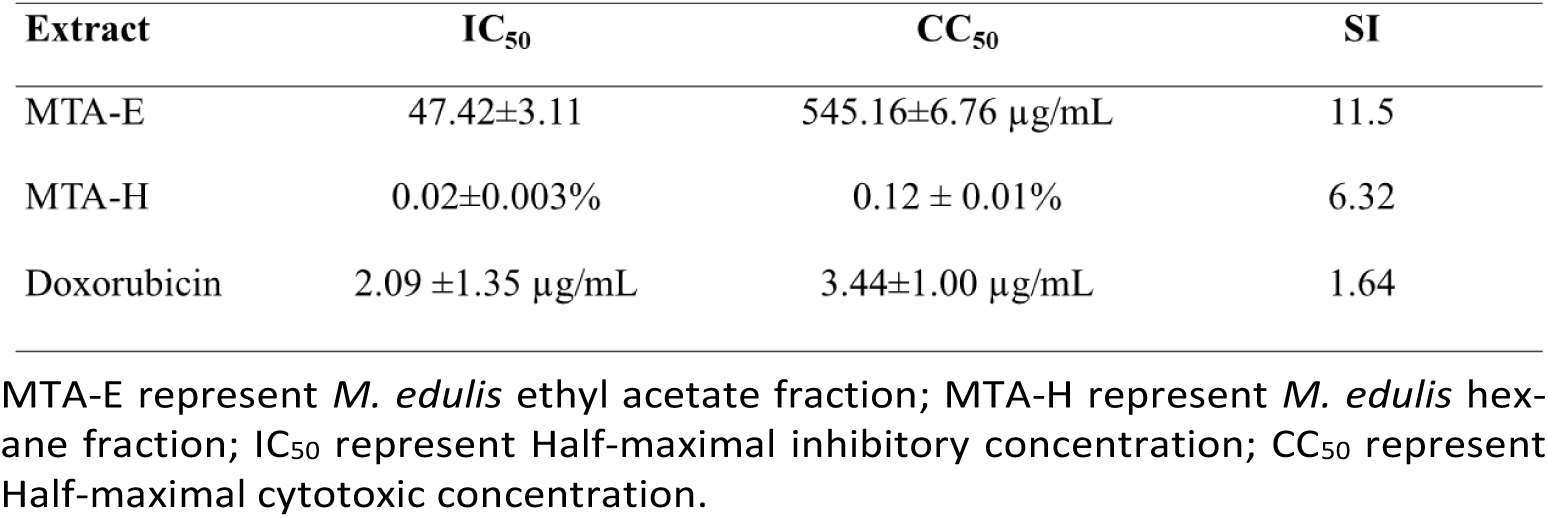
Selectivity index (SI) of *M. edulis* fractions and doxorubicin.

| Extract | IC<sub>50</sub> | CC<sub>50</sub> | SI |
| --- | --- | --- | --- |
| MTA-E | 47.42±3.11 | 545.16±6.76 μg/mL | 11.5 |
| MTA-H | 0.02±0.003% | 0.12 ± 0.01% | 6.32 |
| Doxorubicin | 2.09 ±1.35 μg/mL | 3.44±1.00 μg/mL | 1.64 |

MTA-E represent *M. edulis* ethyl acetate fraction; MTA-H represent *M. edulis* hexane fraction; IC<sub>50</sub> represent Half-maximal inhibitory concentration; CC<sub>50</sub> represent Half-maximal cytotoxic concentration.

**Fig. 4.**
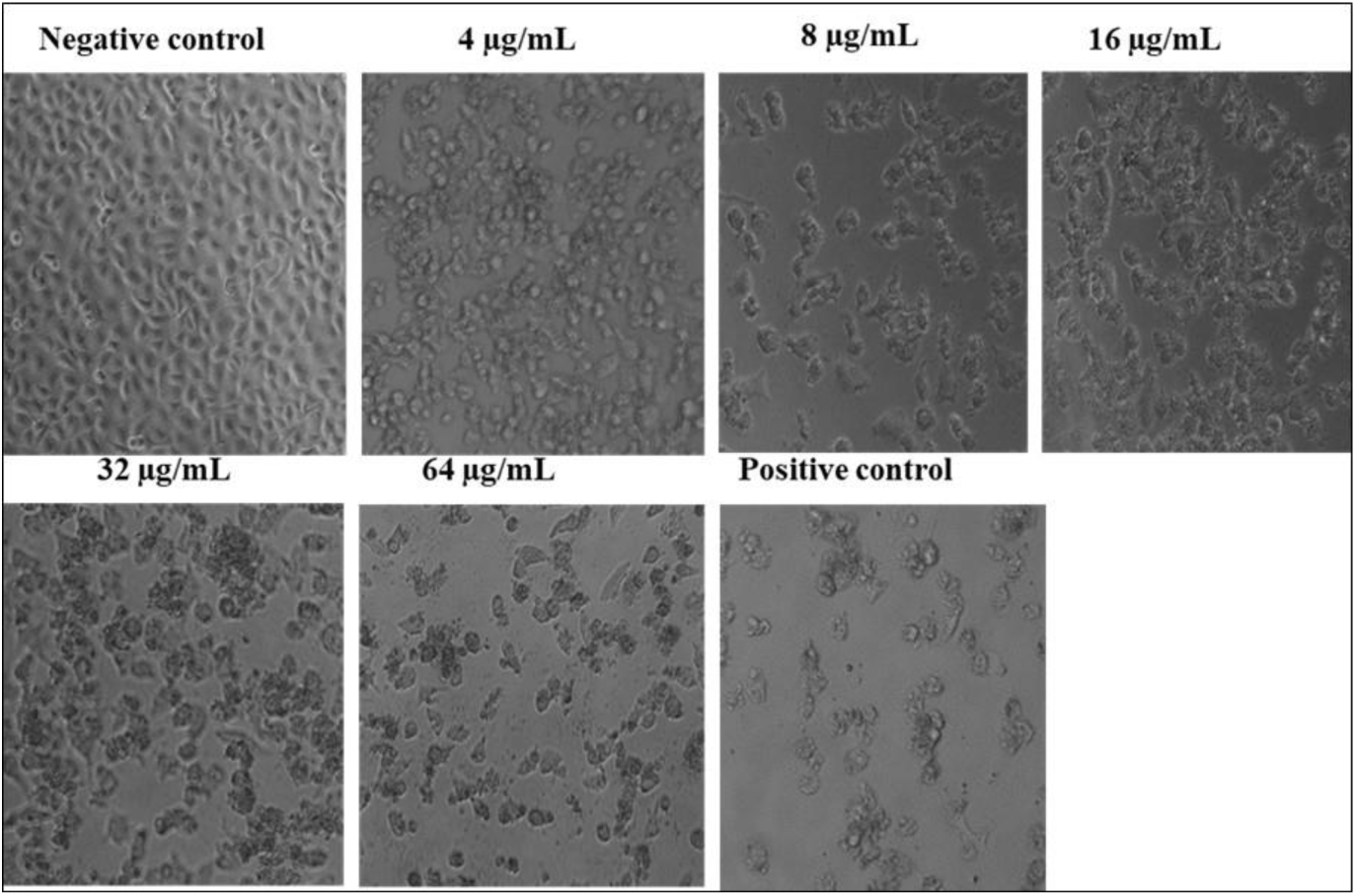
Morphological changes in HeLa cells after 48 hours of exposure to different concentrations of *M. edulis* ethyl acetate fraction. 0.2% DMSO served as the negative control, and doxorubicin as the positive control.

**Fig. 5.**
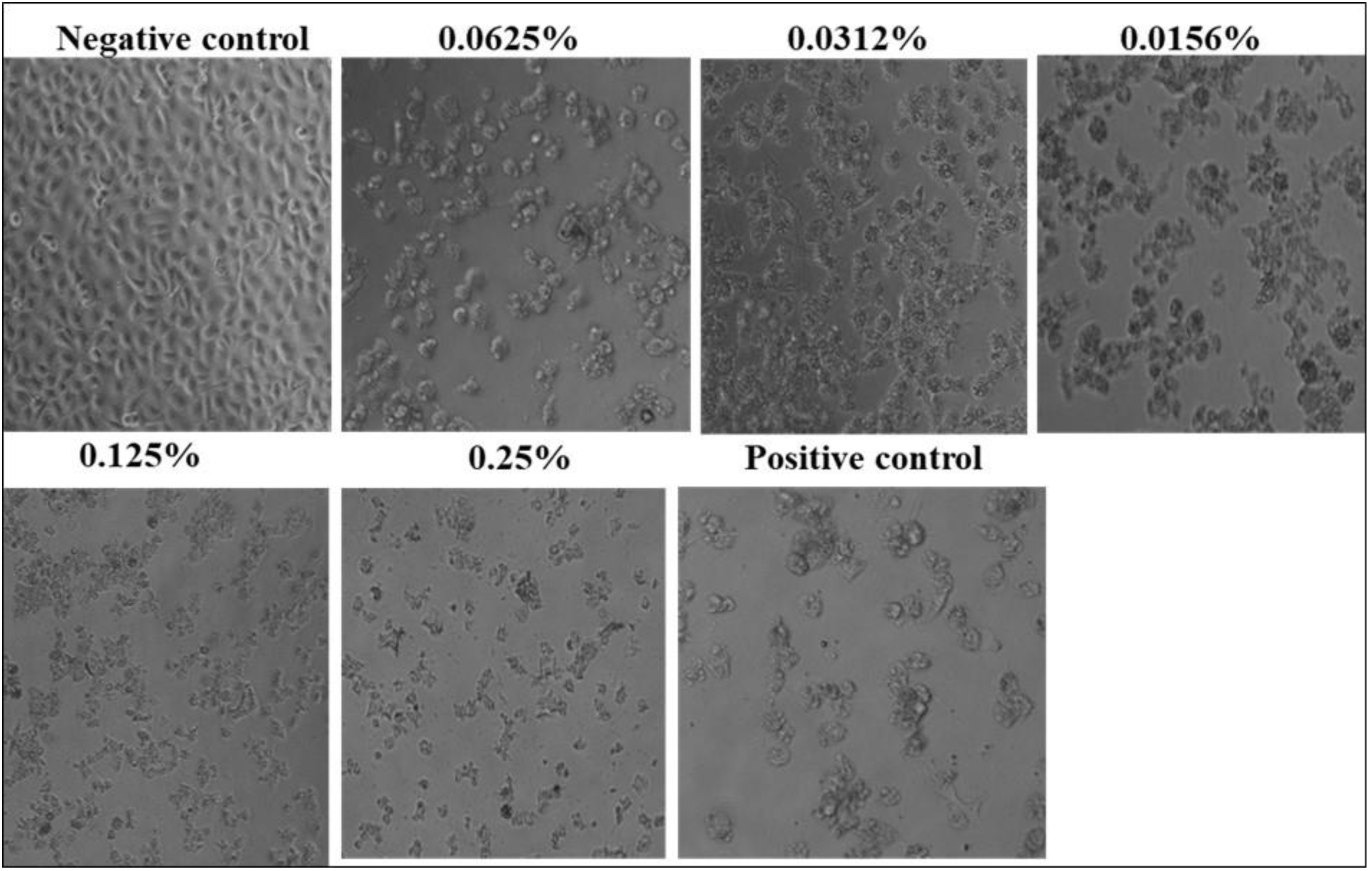
Morphological changes in HeLa cells after 48 hours of exposure to different concentrations of *M. edulis* hexane fraction. 0.2% DMSO served as the negative control, and doxorubicin as the positive control.

### 3.5 Ethyl acetate and hexane fractions of *M. edulis* suppress the migrative ability of HeLa cells

The anti-migratory effects of *M. edulis* ethyl acetate and hexane fractions were evaluated using a wound-healing assay. At the IC_50_ concentrations of 47.24 *µ*g*/*mL (ethyl acetate fraction) and 0.02% (hexane fraction), a significant suppression of HeLa cell migration was observed (Figures 6 and 7). Untreated HeLa cells exhibited normal migration, with complete wound closure after 48 hours. However, cells treated with *M. edulis* fractions demonstrated a reduced wound closure rate (*p <* 0.05). The effect was not time-dependent, as the wound sizes remained significantly larger compared to controls after 24 and 48 hours.

**Fig. 6.**
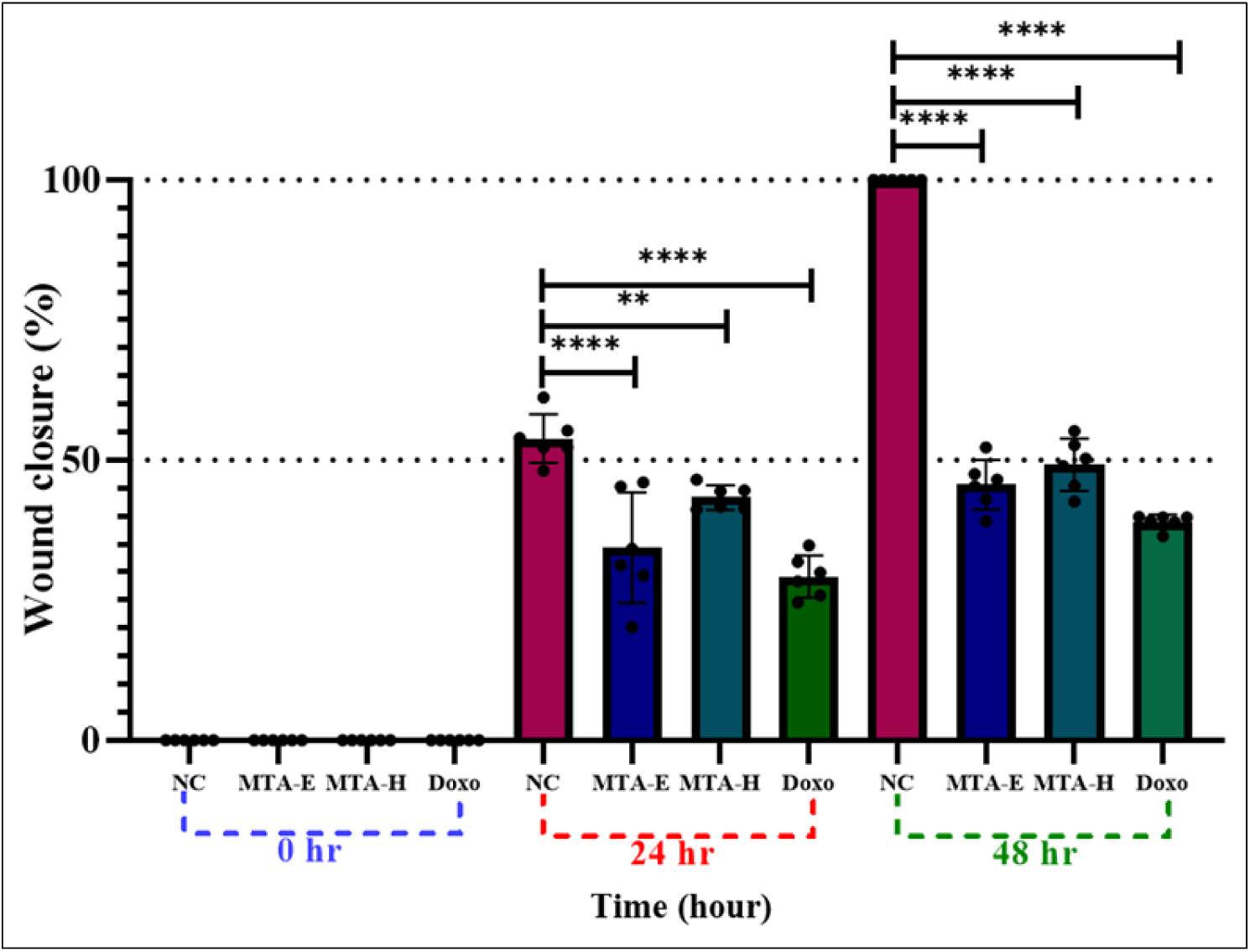
Effect of *M. edulis* ethyl acetate fraction (MTA-E) and hexane fraction (MTA-H) on the migration ability of HeLa cells after 24 and 48 hours of treatment at non-toxic concentrations (*IC*50 of 47.24 *µ*g*/*mL and 0.02%, respectively). The wound closure percentages were significantly (*p <* 0.05) lower compared to the negative control (NC). Data are presented as mean ± SD of five independent experiments performed in triplicate. ** *p <* 0.001, **** *p <* 0.0001.

**Fig. 7.**
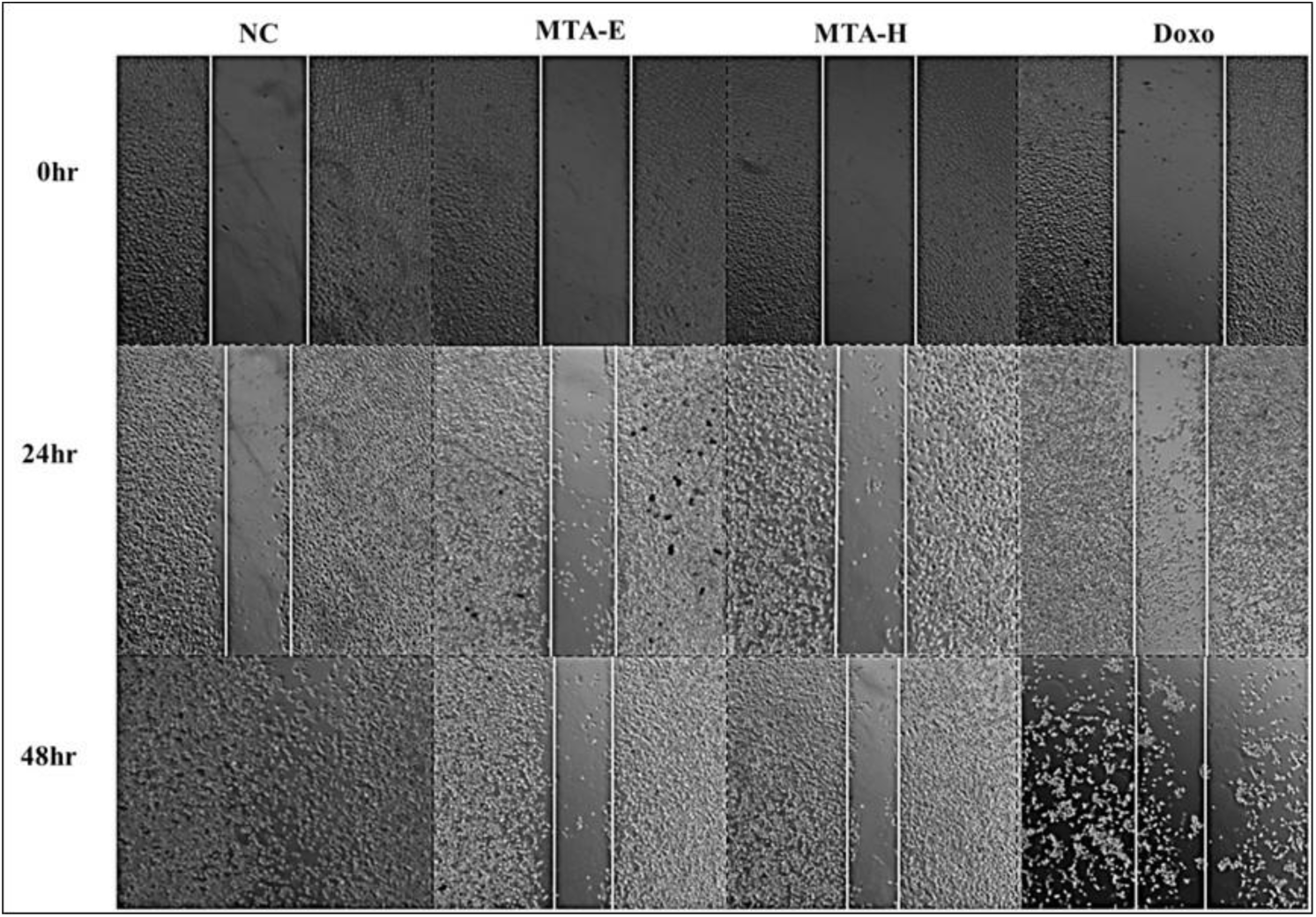
Photomicrographs showing the anti-migratory effects of *M. edulis* ethyl acetate (MTA-E) and hexane (MTA-H) fractions on HeLa cells compared to the negative control (NC) at 0, 24, and 48 hours. A reduction in migration and wound closure was observed over time.

### 3.6 Preliminary screening and GC-MS analysis of *M. edulis* ethyl acetate and hexane fractions

Preliminary phytochemical screening revealed the presence of terpenoids, tannins, and steroids in both ethyl acetate and hexane fractions, while flavonoids were only found in the hexane fraction. Alkaloids, glycosides, phenols, saponins, and quinones were not detected (Table 2). These phytochemicals may contribute to the observed antiproliferation activity. GC-MS analysis identified 33 and 23 peaks in the ethyl acetate and hexane fractions, respectively (Figure 8). Detailed information about the identified phytochemicals, including retention times, molecular weights, and classes, is provided in (Tables 3 and 4).

**Fig. 8.**
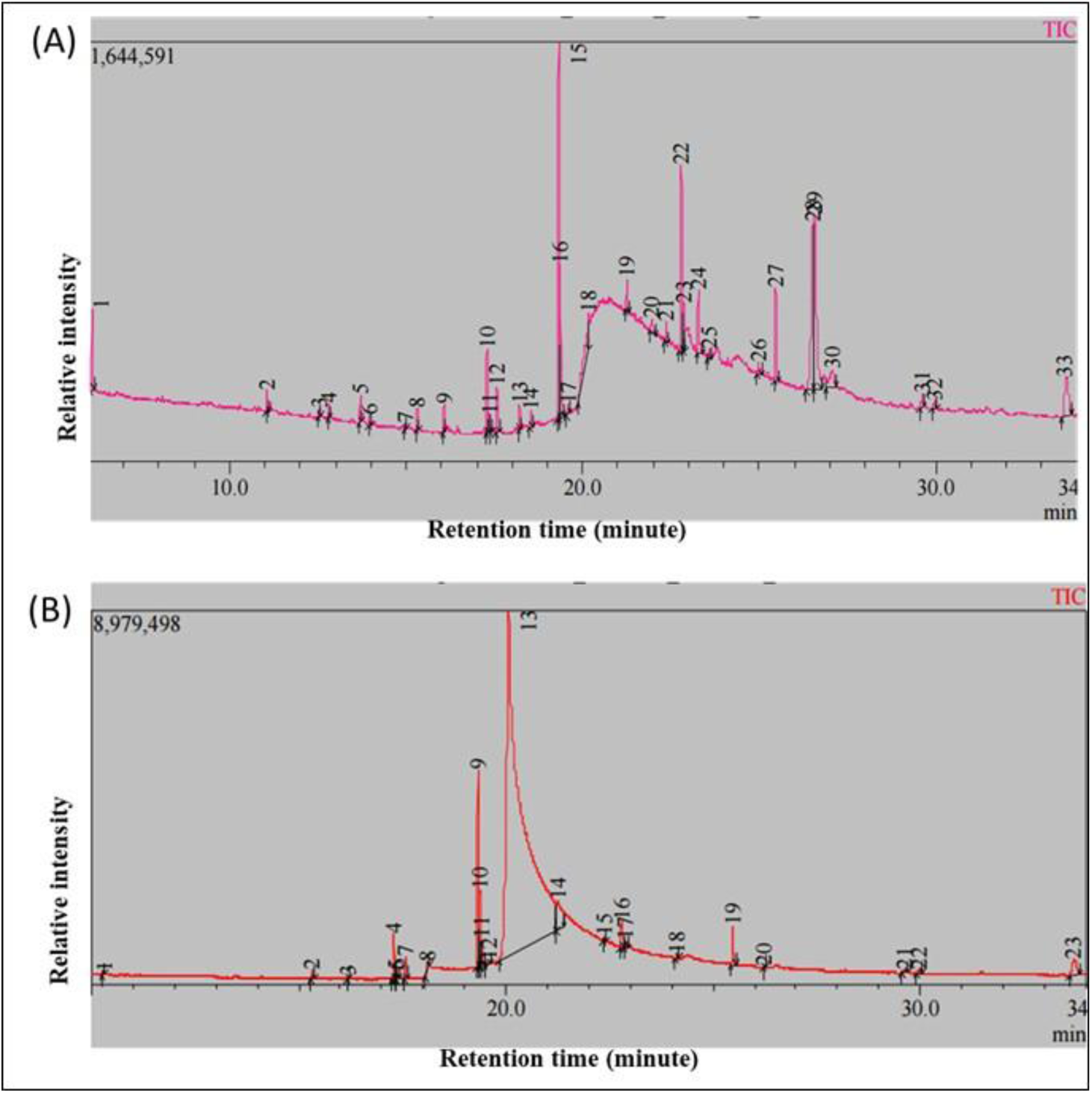
Gas Chromatography-Mass Spectrometry (GC-MS) chromatograms of *M. edulis*. (A) Ethyl acetate fraction; (B) hexane fraction.

**Table 2:** Phytochemical composition of *M. edulis* ethyl acetate (MTA-E) and hexane (MTA-H) fractions.

| Phytochemical | MTA-H | MTA-E |
| --- | --- | --- |
| Alkaloids | - | - |
| Glycosides | - | - |
| Phenols | - | - |
| Flavonoids | + | - |
| Steroids | + | +++ |
| Saponins | - | - |
| Terpenoids | ++ | +++ |
| Quinones | - | - |
| Tannins | ++ | + |

**Key:** - represent absent; + represent low amount; ++ represent moderate amount; +++ represent high amount.

**Table 3:**
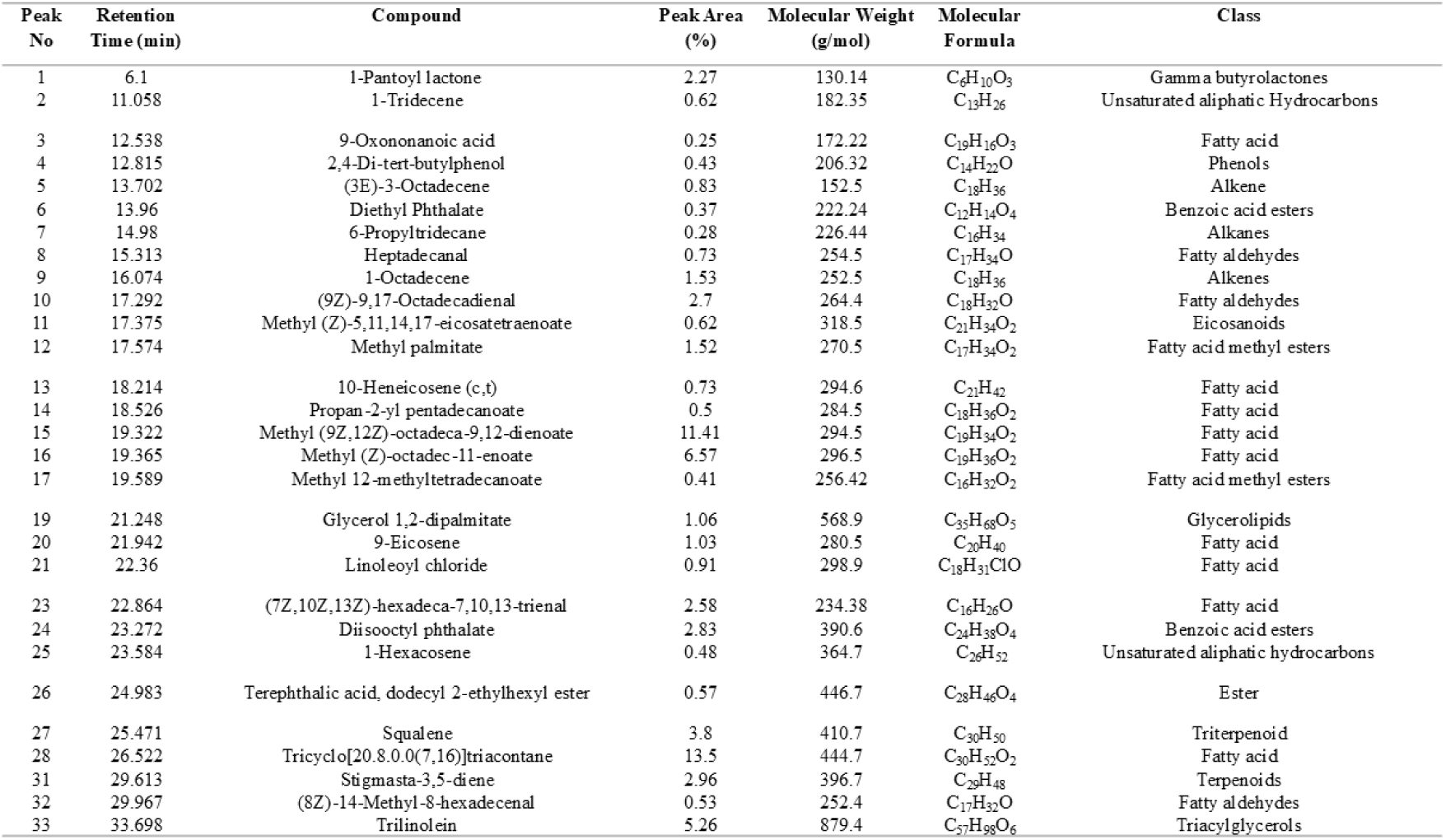
Phytochemicals identified in ethyl acetate fraction of *M. edulis* by GC-MS.

**Table 4:**
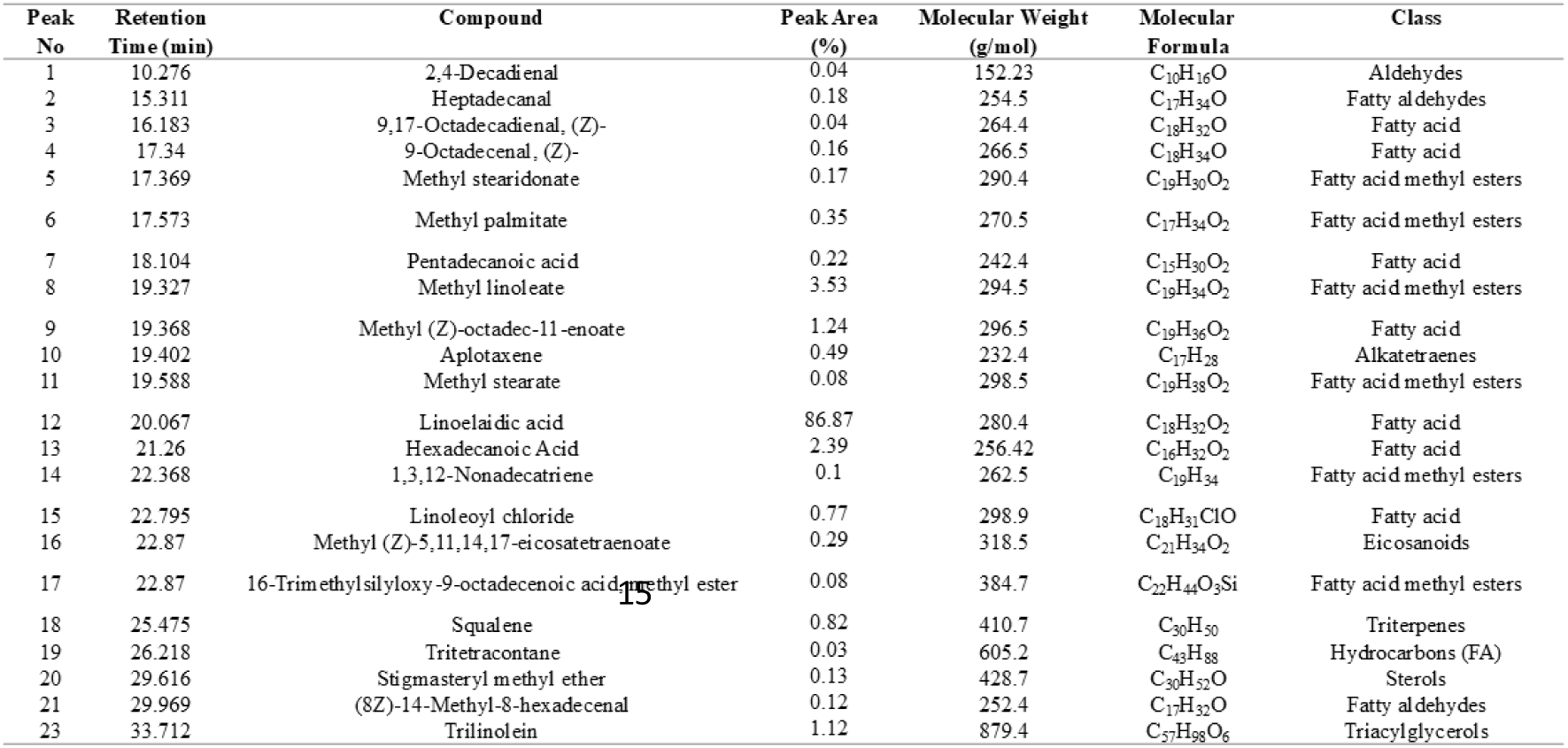
Phytochemicals identified in hexane fraction of *M. edulis* by GC-MS.

## 4 In-silico Results

### 4.1 Screening for Drug-like Phytochemicals in *M. edulis* Ethyl Acetate and Hexane Fractions

The phytochemicals identified in *M. edulis* fractions through GC-MS analysis were subjected to drug-likeness screening. A total of 20 out of 33 and 14 out of 23 phytochemical compounds from the ethyl acetate and hexane fractions, respectively, met Lipinski’s rules (LR) criteria, which include molecular weight (*MW <* 500 g*/*mol), Log P (*<* 5), hydrogen bond acceptors (*HBA <* 10), hydrogen bond donors (*HBD <* 5), and rotatable bonds (*RB <* 10). These compounds were also predicted to have no interference with the blood-brain barrier (BBB) or central nervous system (CNS) (Supplementary Table 2 and 3, respectively). The prioritized phytochemicals were subsequently used for network and molecular docking analyses. Additionally, the ADME-Tox profile prediction suggested that all drug-like phytochemicals exhibited favourable pharmacokinetics properties with minimal potential for associated risks (Supplementary Figures 4 and 5), thus implying a low likelihood of drug-drug interactions. All drug-like (DL) phytochemicals in both fractions demonstrated excellent absorption profiles, with human intestinal absorption (HIA) rates exceeding 90%. The prediction for metabolic properties suggests that the selected phytochemicals do not inhibit cytochromes CYP2D6, CYP2C19, CYP2C9, or CYP3A4, except for minimal inhibition of CYP3A4 and CYP1A2 in both ethyl acetate and hexane fractions of *M. edulis*. ADME-Tox screening further predicted no hepatotoxicity or significant toxicity risks; however, sensitivity to skin upon exposure was predicted. Oral bioavailability was also evaluated using a bioavailability radar (Supplementary Figures 4 and 5). The evaluation was based on six ideally adapted physicochemical profiles for oral drugs, including lipophilicity, polarity, size, solubility, saturation, and flexibility. The thresholds used were as follows:

- Lipophilicity (LIPO): 0.7 *<* XLOGP3 *<* +5
- Size: 150 *<* MV *<* 500 g*/*mol
- Flexibility (FLEX): 0 *<* Number of Rotatable Bonds *<* 9
- Saturation (INSATU): 0.25 *<* Fraction Csp3 *<* 1
- Insolubility (INSOLU): 6 *<* LOG S *<* 0
- Polarity (POLAR): 20 Å^2^ *<* TPSA *<* 130 Å^2^

All drug-like candidates showed favorable oral bioavailability, as evidenced by their placement within the pink zone area of the bioavailability radar. These findings suggest that all drug-like candidates would likely achieve successful absorption and systemic exposure following oral administration.

### 4.2 Active drug-like compounds and cervical cancer-related targets

Potential targets of 20 and 14 phytochemicals of *M. edulis* ethyl acetate and hexane fractions were predicted and retrieved from STP, BDB, and SEA databases. A total of 1926 and 1299 potential targets were identified for ethyl acetate and hexane fractions, respectively. After the removal of duplicated targets from the pools in each fraction, 811 and 642 unique potential targets were obtained. Further, a total of 5809 cervical cancer-related genes were retrieved from the GeneCards, DisGeNET, Pharos, and OMIM databases. After removal of duplicated targets, 2953 unique targets were obtained. All targets were standardized using the UniProt database. A Venn diagram was generated to show the intersected targets between the *M. edulis* ethyl acetate and hexane fractions and cervical cancer targets. A total of 211 and 182 overlapping potential targets were obtained, respectively (Figure 9A-B and Extended Supplementary Table 1 and 2)

**Fig. 9.**
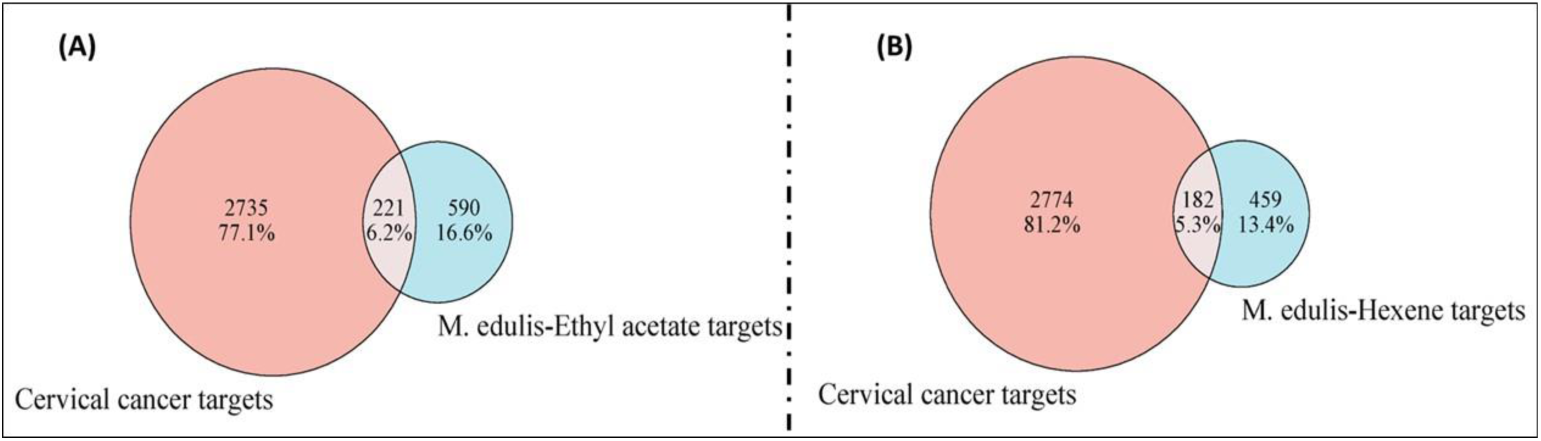
Venn diagram of the intersected targets between M. edulis’ fractions phytochemicals and cervical cancer. (A) M. edulis ethyl acetate-cervical cancer targets, (B) M. edulis hexane-cervical cancer targets

### 4.3 Protein-protein interactions (PPI) network map and the key targets

A total of 211 and 182 target genes for ethyl acetate and hexane fractions of *M. edulis* were imported into the STRING database to construct PPI network maps. Using a high confidence interaction score of *>* 0.6, the PPI network of *M. edulis* ethyl acetate and hexane therapy for cervical cancer was generated. The ethyl acetate fraction network exhibited; 211 nodes and 3707 edges (expected edges of 1618), Average node degree of 33.5, Average local clustering coefficient of 0.527, PPI enrichment p-value of *<* 0.01*e* − 16 (Figure 10A). The hexane fraction network yielded 182 nodes and 2624 edges (expected edges of 1134), Average node degree of 28.8, Average local clustering coefficient of 0.577, PPI enrichment p-value of *<* 0.01*e* − 16 (Figure 10B). Cytoscape software v3.8.2 was used to analyze network topological properties. The top 30 hub targets were identified based on their degree scores (degree ≥ 35) for both ethyl acetate and hexane fractions (Figure 11). Notably, 21 common targets were identified among the top 30 hub targets for both fractions, including ABL1, CDK1, CDK2, CDKNIA, EGFR, ERBB2, ESR1, GSK3B, HDAC1, HIF1A, MAPK14, MCL1, MMP9, MTOR, PIK3CA, PPARA, PPARG, PTGS2, SRC, and STAT3 (Supplementary Figure 6). These data suggest that both candidate compounds in ethyl acetate and hexane fractions of *M. edulis* probably acted on the same targets in cervical cancer.

**Fig. 10.**
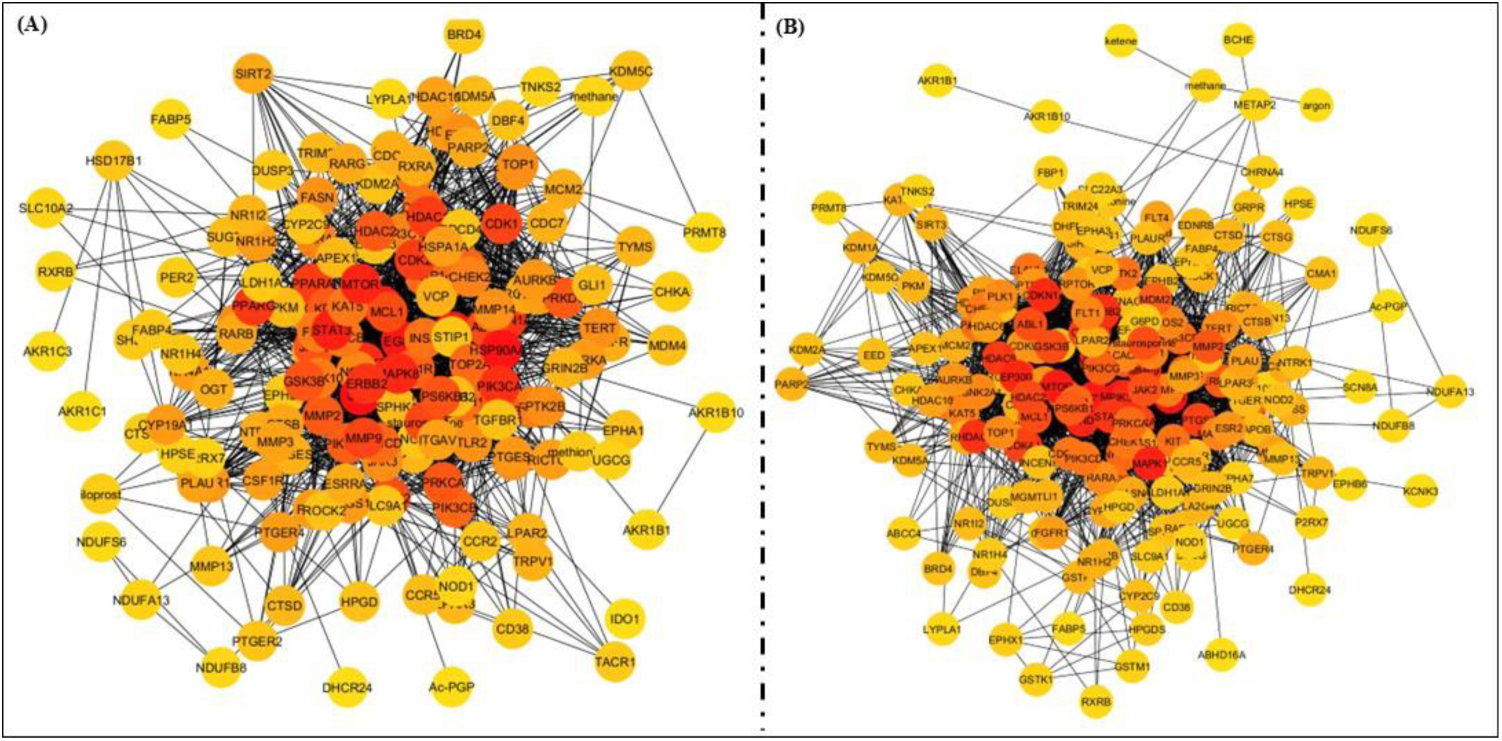
PPI network for *M. edulis* anti-cervical cancer targets. (A) Ethyl acetate and (B) Hexane fractions. Red color intensity indicates higher associations; yellow indicates lower associations.

**Fig. 11.**
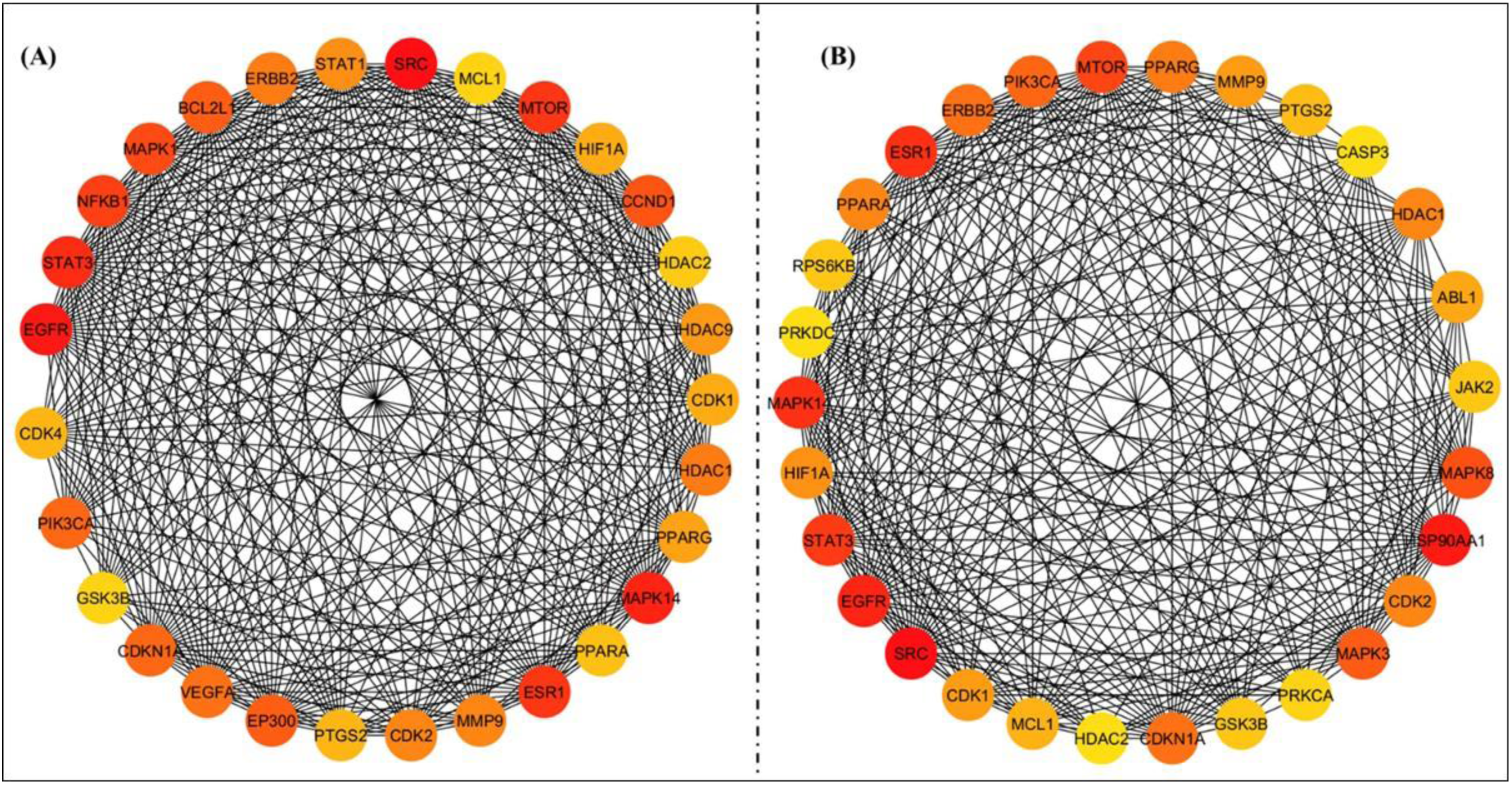
Top 30 hub genes PPI network. (A) Ethyl acetate and (B) Hexane fraction targets. Red color intensity indicates higher association and significance of the target.

### 4.4 Gene ontology (GO) enrichment analysis

GO enrichment analyses were performed on 211 targets of the ethyl acetate fraction and 182 targets of the hexane fraction to unravel the most enriched biological functions. GO terms were classified into three categories:

- **Biological Processes (BP),** whereby, Ethyl acetate had 1817 significant terms, while hexane had 1599 terms, including responses to oxygen-containing compounds, organic substances and regulation of biological quality.
- **Molecular Functions (MF),** whereby, Ethyl acetate had 193 terms, while hexane had 170 terms, including signaling receptor activity, molecular transducer activity, and protein kinase activity.
- **Cellular Components (CC),** whereby, Ethyl acetate had 108 terms, while hexane had 95 terms, focusing on intrinsic and integral components of plasma membranes and nuclear lumens.

The top 15 GO terms significantly enriched are presented in Figure 12A-C and 12X-Z for ethyl acetate and hexane, respectively.

**Fig. 12.**
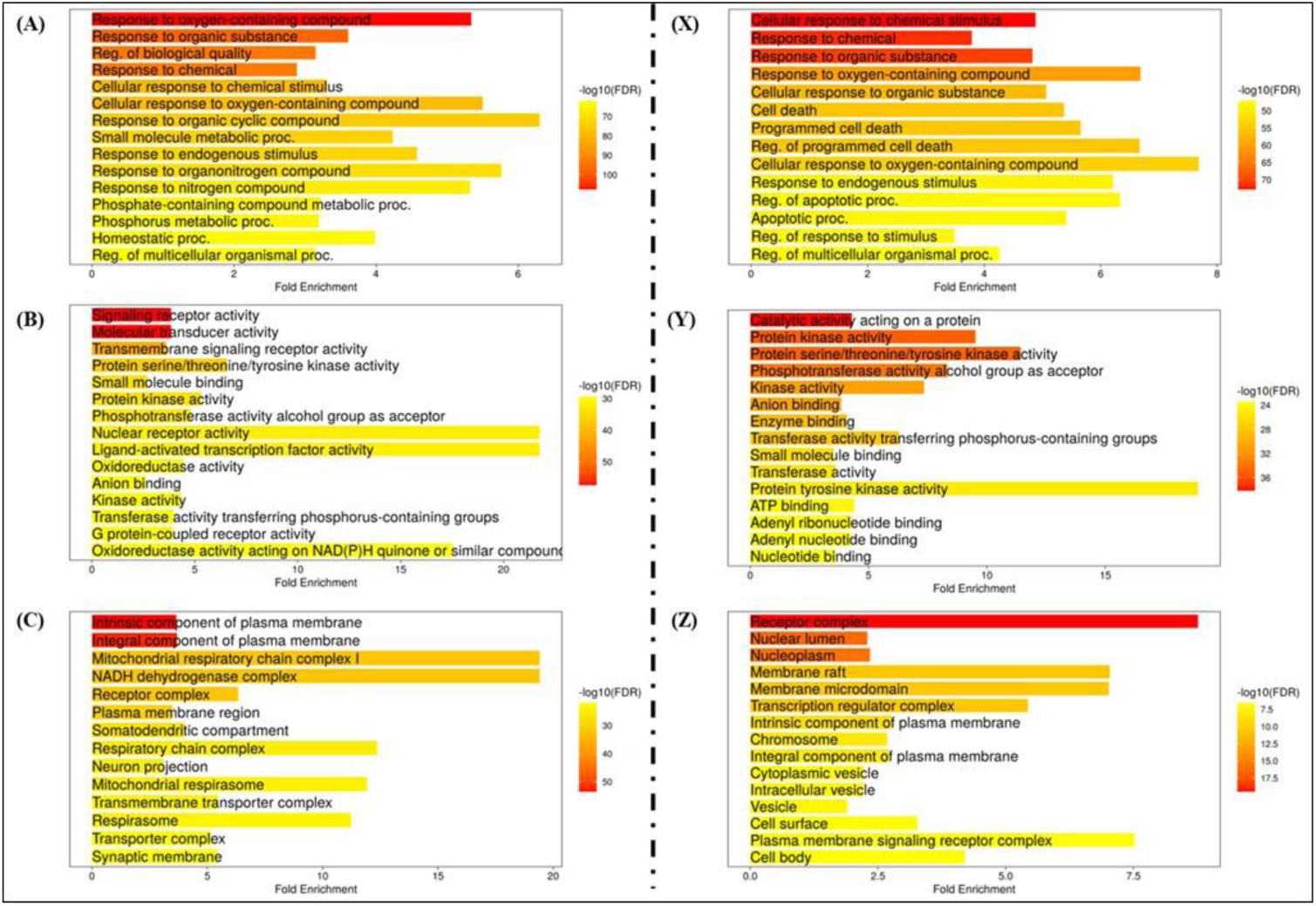
GO enrichment analysis for the 221 and 182 anticervical cancer targets of ethyl acetate and hexane of M. edulis. A-C presents GO terms of ethyl acetate and X-Z presents GO terms of hexane fractions. The X-axis presents the number of target fold enrichments; the Y-axis shows AX; biological process (AP), BY; cellular components (CC), and CZ; molecular function (MF). Bar enclosing GO terms with red colour indicates the significance of the enriched terms

### 4.5 Kyoto Encyclopedia of Genes and Genomes (KEGG) Pathway Enrichment Analyses

KEGG pathway enrichment analysis was performed to understand better the molecular therapeutic mode of action of *M. edulis* fractions target genes. Interestingly, 184 and 178 putative pathways were significantly enriched (P *<*0.05) for ethyl acetate and hexane fractions, respectively. Ethyl acetate targets are mainly enriched with the following pathways; pathway in cancer, proteoglycan in cancer, PI3K-Akt1 signaling pathway and microRNAs in cancer pathways (Figure 13A). Hexane extract’s top enriched pathways included neuroactive ligand-receptor interaction, metabolic pathways, retrograde endocannabinoid signalling and pathways (Figure 13B). The top 15 enriched pathways are presented in the barplot (Figure 13A-B). The red colour intensity signifies the lowest p-value and more significantly enriched pathways. To further elucidate the putative mechanisms of *M. edulis* fraction for potential cervical cancer treatment, we constructed a functional mapping classifying significant KEGG pathways (Figure 14A-B).

**Fig. 13.**
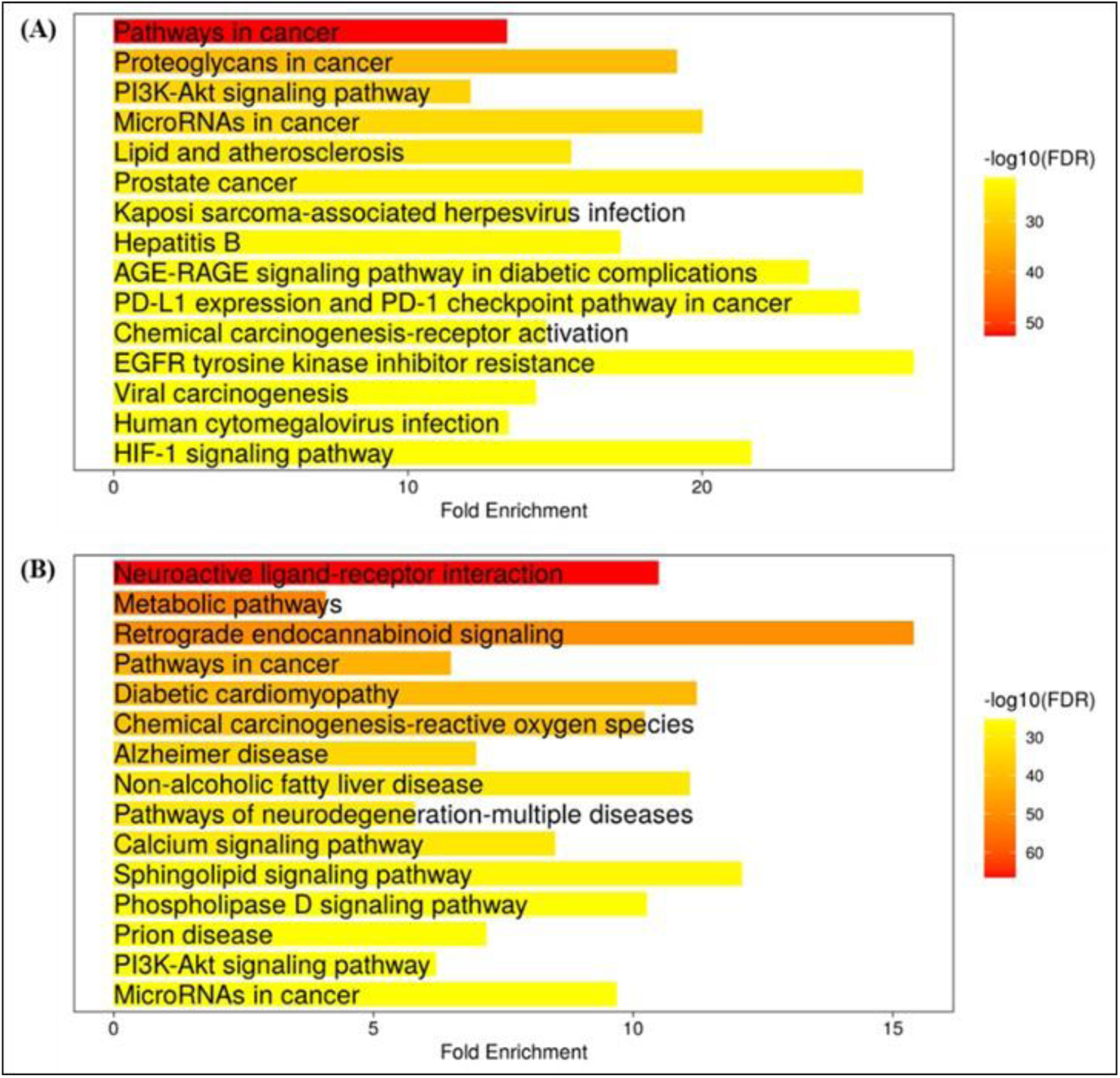
KEGG pathway enrichment analysis for the 221 and 182 anticervical cancer targets of ethyl acetate and hexane fractions of M. edulis. The X-axis displays the number of target fold enrichments; the Y-axis show the various KEGG pathways. Bar enclosing pathways with red colour indicates the significance of the enriched pathways

**Fig. 14.**
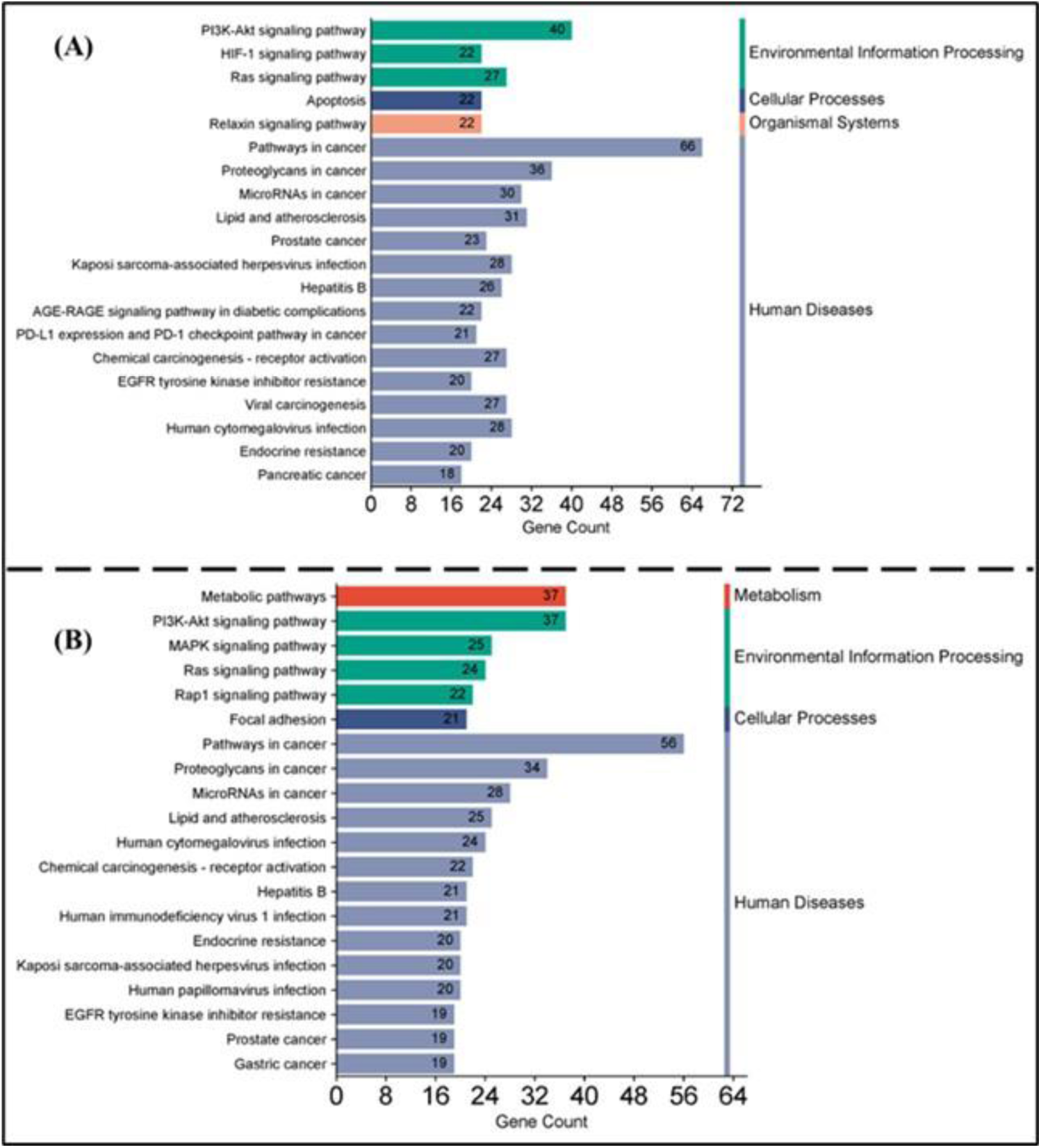
KEGG pathway mapping of cervical cancer-related targets. (A) Ethyl acetate and (B) Hexane fractions of *M. edulis*.

### 4.6 Molecular Docking Results

A molecular docking study was conducted to validate the targets and investigate the putative mechanistic role of individual phytochemicals of *M. edulis* toward anticervical potential. The docking results indicated that ligands such as diisooctyl phthalate (E15), squalene (E18 and H12), stigmasta-3,5-diene (E20), (8Z)-14-methylhexadec-8-enal (E21 and H14), methyl stearate (H7), and methyl-5,11,14,17-eicosatetraenoate (H10) had great putative binding affinity for selected cervical cancer-related proteins, namely BCL2, CDK2, CDK1, TP53, CASP9, P21, HDAC1, and CCDN1. Various binding affinity scores shown in (Table 6) demonstrate how well the selected proteins and ligands can interact. The affinity score obtained from the docking results ranges from −3.5 to −8.5 kcal/mol. Affinity value indicates the strength of how a ligand interacts with a protein target. However, the lower the binding affinity, the stronger the binding interaction between the molecules [80]. The docking results were validated by redocking the native ligands against the selected targeted proteins (Supplementary Figure 7 and Supplementary Table 4). All the ligands analysed showing binding affinity to the receptors below −7.0 kcal/mol are presented in 2D models for ethyl acetate fraction (Figure 15). E15 compound binds onto the receptor-binding pocket of chain B of BCL2 protein (−7.7 kcal/mol) specifically on the following residues: Val115, Glu119, Val118, Lys22, Phe71, asp61, Arg57, Tyr18, Val60, Tyr21 with the following bonds; van der Waals, conventional hydrogen bond, alkyl and Pi-alkyl (Figure 15A) whereas, NL-1 binds onto the receptor-pocket of chains A and C of BCL2 protein (−8.7 kcal/mol) through the following residues: Arg26, Asp61, Lys22, Glu25, Phe71, Val118, Val115, Gln119 to form conventional hydrogen bond, P-cation, Pi-anion, Pi-sigma and Pi-alkyl interactions (Supplementary Figure 7A). E15 compound binds onto the receptor-binding pocket of chain A of CDK2 protein (−8.3 kcal/mol) specifically on the following residues: Tyr15, Ala151, Ile52, Ile35, Phe152, Leu78, Leu148, Leu55, Leu66, Phe80 with the following bonds: alkyl and Pi alkyl (Figure 11B). E20 compound binds onto the receptor-binding pocket of chain A of CDK2 protein (−8.1 kcal/mol) specifically on the following residues: Tyr180, Arg169, Asp127, Gly147, Thr14, Lys33, Ala151, Phe152, Ile35, Phe146, Leu78, Ile52, Leu148 through van de Waals and alkyl (Figure 15C) whereas NL-2 binds onto the receptor-binding pocket of chain A of CDK2 protein (−8.7 kcal/mol) through the following residues: Lys129, Thr14, Lys33, Asp127, Leu78, Ile52, Phe35, Phe152, Ala151, Leu148 through carbon hydrogen bond, P-cation, Pi-anion, alkyl, P-alkyl and unfavourable donor-donor interactions (Supplementary Figure 7B). E20 compound binds onto the receptor-binding pocket of chain B of CDK1 protein (−7.4 kcal/mol) specifically on the following residues: Tyr367, Val170, Ile311, Lys310 through alkyl and Pi alkyl bonds (Figure 15D) whereas NL-3 binds onto the receptor-pocket of chain A of CDK1 protein (−7.6 kcal/mol) through the following residues: Thr15, Lz9301, Val165, Gln132, Glu12 forming conventional hydrogen bond, carbon hydrogen bond, Pi-donor hydrogen bond, Pi-sigma, alkyl and Pi-alkyl interactions (Supplementary Figure 7C). E18 compound binds onto the receptor-binding pocket of chain A of TP53 protein (−7.9 kcal/mol) specifically on the following residues: Arg99, Ser96, Glu97, Lys95, Val98, Val164, Leu65, Gln165, Val163, Asp63, Pro94 through van de Waals and alkyl bonds (Figure 15E), whereas NL-4 binds onto the receptor-pocket of chain A of TP53 protein (−5.2 kcal/mol) through the following residues: Pro94, Lys95, Leu65, Val98, Val163, Glu97, Val164, forming conventional hydrogen bond, Pi-sigma, alkyl and Pi-alkyl interactions (Supplementary Figure 7D). E15 binds onto the receptor-binding pocket of chains A and B of P21 protein (−7.5 kcal/mol) specifically on the following residues: Pro307, Lys308, Val425, Ile312, Pro369, Val425, Leu470 through carbon hydrogen bond and alkyl (Figure 11F). On the other hand, E20 binds onto the receptor-binding pocket of chains A and B of P21 protein (−8.1 kcal/mol) specifically on the following residues: Pro307, Val425, Lys308, Leu311, Phe410, Pro469, Met424 using conventional hydrogen bond, alkyl and Pi-alkyl Pi alkyl bond (Figure 15G) whereas NL-6 binds onto the receptor-pocket of chains A and B of P21 protein (−8.5 kcal/mol) through the following residues: Phe410, Leu311, Gly409, Ile312, Asp407, Gln423, Asp393, forming convectional hydrogen bond, carbon hydrogen bond, Pi-anion, Pi-Pi T-shaped, alkyl and Pi-alkyl interactions (Supplementary Figure 7F). E15 compound binds onto the receptor-binding pocket of chains A and B of HDAC1 protein (−8.1 kcal/mol) specifically on the following residues: Ile249, Asp332, Met329, Arg36, Tyr333, Asn40, Asp256, Asp248, Tyr15, His19, Tyr48, Phe252 through the following bonds: van de Waals, conventional hydrogen bond and Pi-alkyl (Figure 15H) whereas E20 compound binds onto the receptor-binding pocket of chains A and B of HDAC1 protein (−8.5 kcal/mol) specifically on the following residues: Asp248, Asn40, Arg36, His39, Tyr15, Gly17, His259, Lys260, Phe252, Ile249, Arg55, Val198, Gln197, Thr196, Asp256 through the van de Waals, alkyl and Pi-alkyl bonds (Figure 15I). NL-7 binds onto the receptor-pocket of chains A and B of HDAC1 protein (−5.5 kcal/mol) through the following residues: Phe252, Asp256, Glu325, Arg36, Asn21, Asp18, Asp16, Gly17 forming conventional hydrogen bond, carbon hydrogen bond, Pi-cation, Pi-donor hydrogen bond as well as attractive charge and unfavourable positive-positive interactions (Supplementary Figure 7G). E15 compound binds onto the receptor-binding pocket of chain B of CCND1 protein (−7.3 kcal/mol) specifically on the following residues: Asp96, Glu141, Val93, Asp94, Leu144, Lys35, Asp155, Leu158, Phe90, Phe156, Leu470, Ala154, Glu91, Val69, His92, Asn142 through the van de Waals, alkyl and Pi-alkyl bonds (Figure 15J), whereas NL-8 binds onto the receptor-pocket of chains A and B of CCND1 protein (−4.2 kcal/mol) through the following residues: Gly127, Ser54, Ser33 to form conventional hydrogen bond, carbon hydrogen bond and unfavourable donor-donor interactions (Supplementary Figure 7H).

**Fig. 15.**
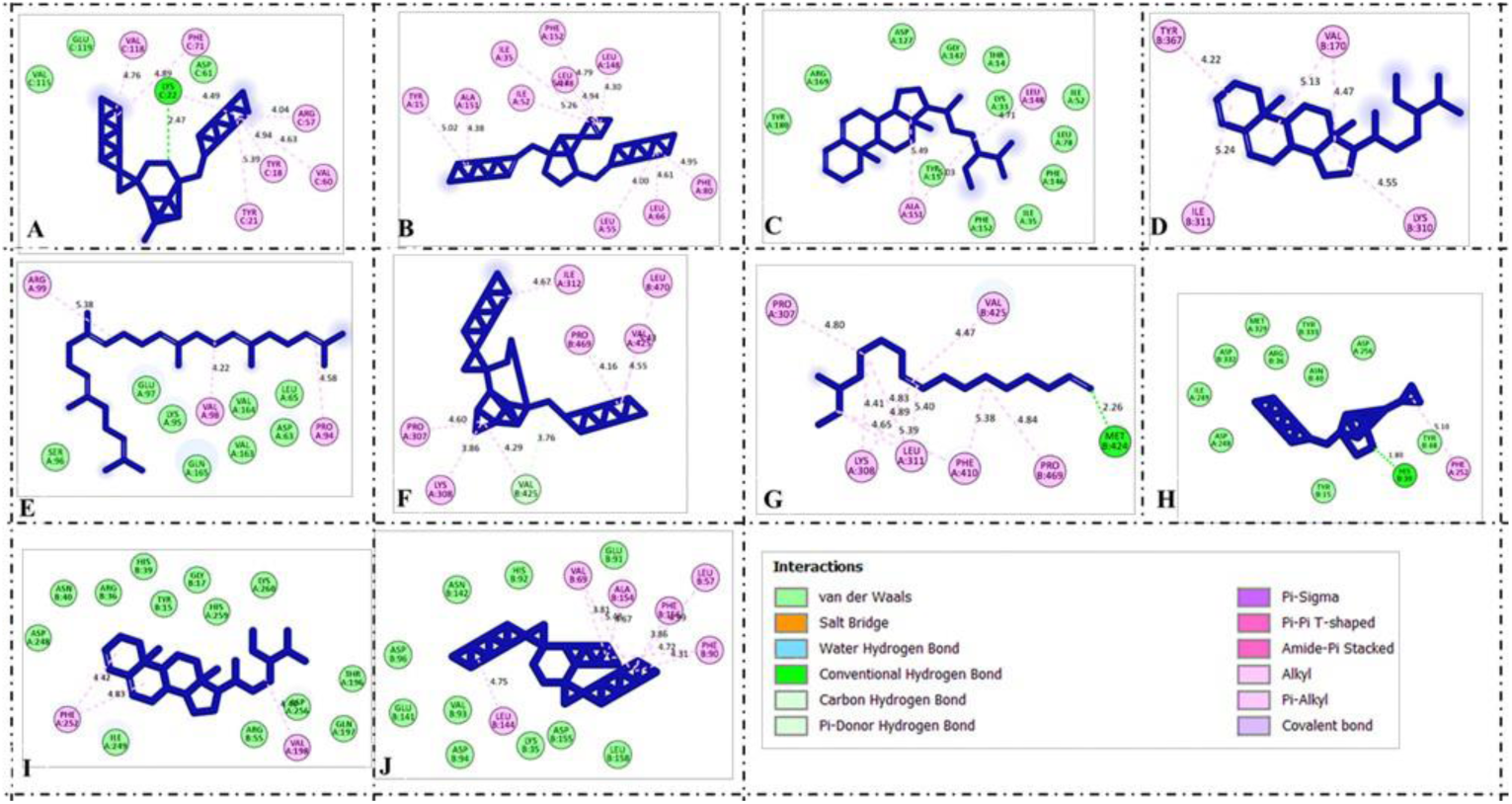
Two-dimension (2D) molecular docking results. Binding modes of compounds of ethyl acetate fraction onto binding-receptor pockets of the selected target protein. A) Diisooctyl phthalate bind BCL2, B) & C) Diisooctyl phthalate and stigmasta-3,5-diene bind CDK2, D) Stigmasta-3,5-diene bind CDK1, E) Squalene bind TP53, F) & G) Diisooctyl phthalate and stigmasta-3,5-diene bind P21, H) & I) Diisooctyl phthalate and stigmasta-3,5-diene bind HDAC1 and J) Diisooctyl phthalate bind CCND1.

**Table 5:**
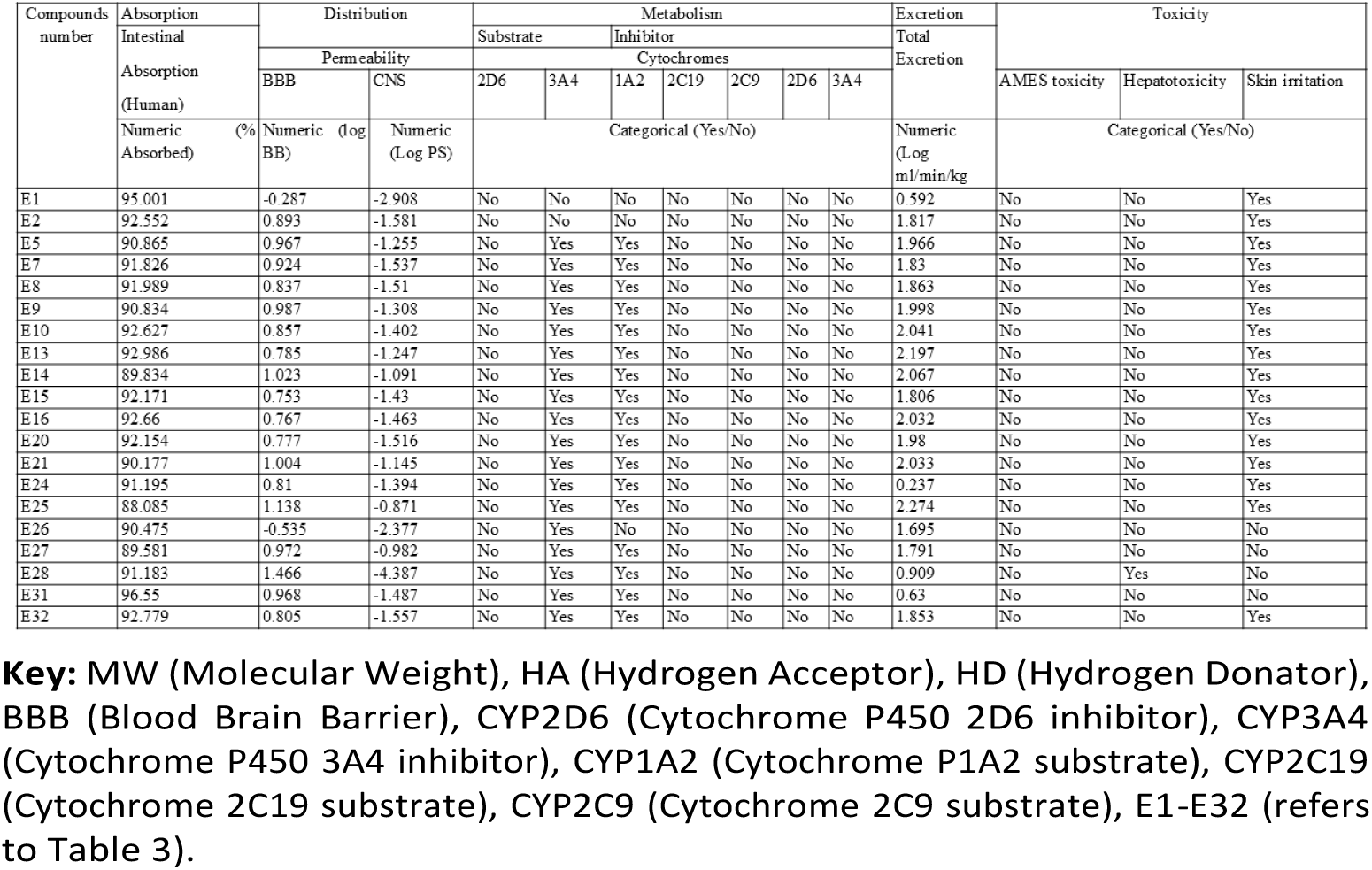
Prediction of ADME-Tox in-silico pharmacokinetic profiles of drug-like phytochemicals in *M. edulis* ethyl acetate fraction.

**Table 6:**
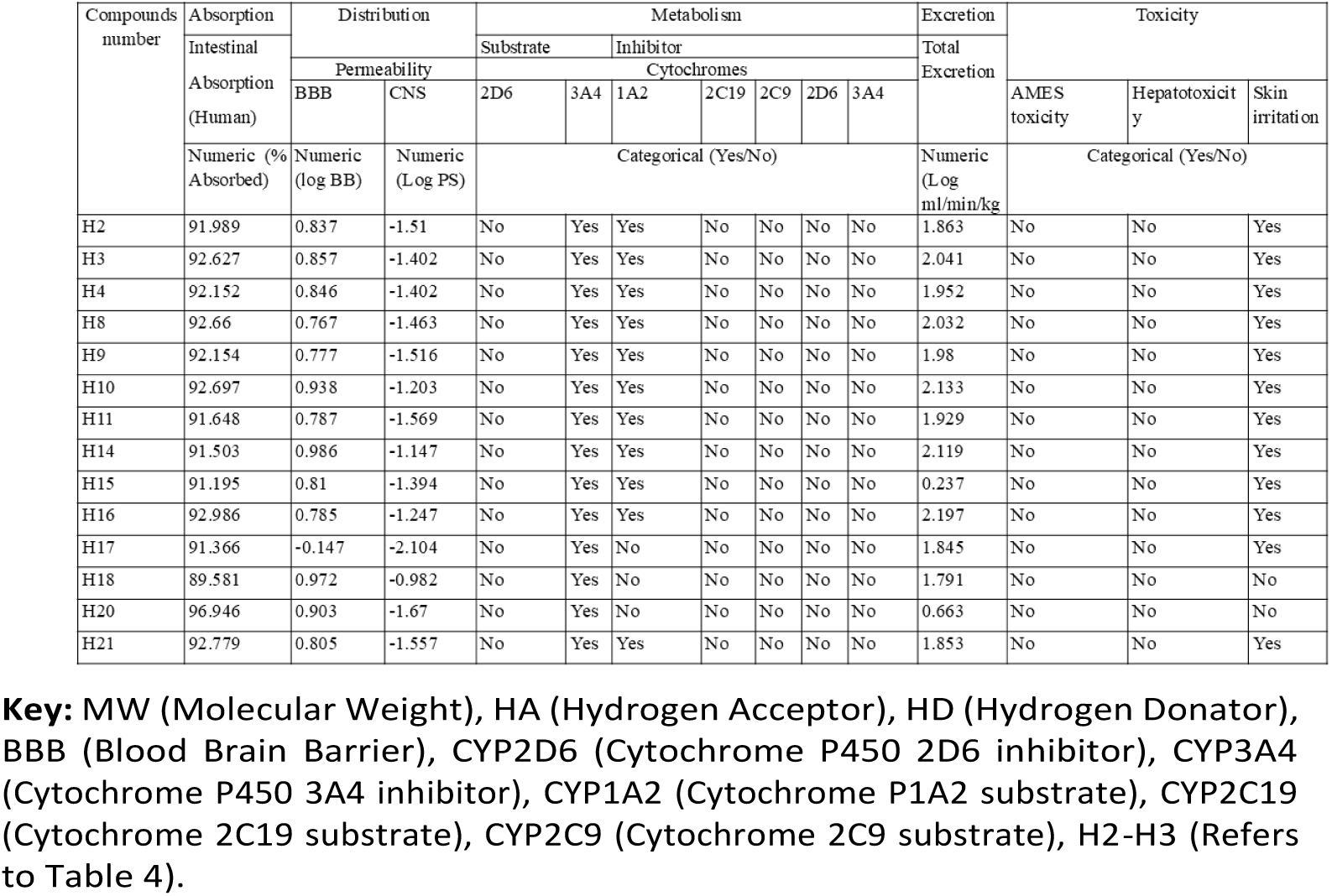
Prediction of ADME-Tox in-silico pharmacokinetic profiles of phytochemicals in *M. edulis* hexane fraction.

**Table 7:** Binding affinity of *M. edulis* compounds with selected target proteins.

| Receptor/<br>Ligand | BCL2 | CDK2 | CDK1 | TP53 | CASP9 | P21 | HDAC1 | CCND1 |
| --- | --- | --- | --- | --- | --- | --- | --- | --- |
| **Ethyl acetate fraction** |  |  |  |  |  |  |  |  |
| **E15** | -7.7 | -8.3 | -7.5 | -6.2 | -6.9 | -7.5 | -8.1 | -7.3 |
| **E18** | -5.7 | -6.9 | -6.9 | -7.9 | -5.8 | -6.3 | -6.5 | -5.9 |
| **E20** | -6.9 | -8.1 | -7.4 | -6.3 | -8.0 | -8.1 | -8.5 | -7.4 |
| **E21** | -4.9 | -7.2 | -4.7 | -3.5 | 4.8 | -5.3 | -5.7 | -4.2 |
| **Hexane fraction** |  |  |  |  |  |  |  |  |
| **H7** | -6.1 | -7.4 | -6.6 | -4.3 | -5.3 | -6.0 | -6.8 | -6.0 |
| **H10** | -4.9 | -6.3 | -5.0 | -3.6 | -4.5 | -5.9 | -5.2 | -4.5 |
| **H12** | -5.5 | -6.9 | -5.5 | -4.2 | -5.1 | -7.0 | -6.1 | -5.1 |
| **H14** | -5.1 | -6.9 | -5.1 | -3.6 | -3.9 | -5.6 | -5.6 | -4.9 |

**Key:** E15-Diisooctyl phthalate; E18-Squalene; E20-Stigmasta-3,5-diene; E21-(8Z)-14-Methylhexadec-8-enal; H7-Methyl stearate; H10-Methyl-5,11,14,17-eicosatetraenoate; H12-Squalene and H14-(8Z)-14-Methylhexadec-8-enal.

All the ligands analysed showing binding affinity to the receptors below −6.0 kcal/mol are presented in 2D models for hexane fraction (Figure 16). H7 compound binds onto the receptor-binding pocket of chain C of BCL2 protein (−6.1 kcal/mol) specifically on the following residues: Tyr139, Val93, Phe89, Ala90, His143, Arg142 through conventional hydrogen bond, carbon hydrogen bond, alkyl and Pi-alkyl bonds (Figure 16A). H7 compound binds onto the receptor-binding pocket of chain A of CDK2 protein (−7.4 kcal/mol) specifically on the following residues: Lys33, Leu66, Leu78, Leu55. Ieu52, Phe80, Ile63, Phe146, Leu58 through the conventional hydrogen bond, alkyl and Pi-alkyl bonds (Figure 16B). H12 compound binds onto the receptor-binding pocket of chain A of CDK2 protein (−6,9 kcal/mol) specifically on the following residues: Ala151, Agr126, Tyr15, Leu148, Phe80, Leu66, Leu66, Leu78, Leu55, Ile52 through the alkyl and Pi-alkyl bonds (Figure 16C). H14 compound binds onto the receptor-binding pocket of chain A of CDK2 protein (−6.9 kcal/mol), specifically on the following residues: Phe146, Leu55, Phe80, Val64, Leu148, Leu66, Ile52, Leu78, Phe152, Ile35 through the alkyl and Pi-alkyl bonds (Figure 16D). H7 compound binds onto the receptor-binding pocket of chain B of CDK1 protein (−6.6 kcal/mol) specifically on the following residues: Phe338, Leu300, Pro299, Val366, Pro301, Tyr223, Met330, Ile343, Val186, Gln184, Ser227, Val226, Thr329 through the van der Waals, carbon hydrogen bond and alkyl bond (Figure 16E). H12 compound binds onto the receptor-binding pocket of chains A and B of P21 protein (−7.4 kcal/mol) specifically on the following residues: Ala280, Phe410, Val425, Leu311, Pro307, Lys308, Val425 through the alkyl and Pi-alkyl bonds (Figure 16F). H7 compound binds onto the receptor-binding pocket of chain B of HDAC1 protein (−7.0 kcal/mol) specifically on the following residues: Phe205, His141, Zn401, His410 through the conventional hydrogen bond, metal-acceptor and alkyl bond (Figure 16G). H7 compound binds onto the receptor-binding pocket of chain A of CCND1 protein (−6.8 kcal/mol) specifically on the following residues: Leu28, His163, Lys167, Val236, Phe232, Arg235 through the conventional hydrogen bonds, alkyl and Pi-alkyl bonds (Figure 16H). We validated our results by redocking using native ligands, as depicted in (Supplementary Figure 7).

**Fig. 16.**
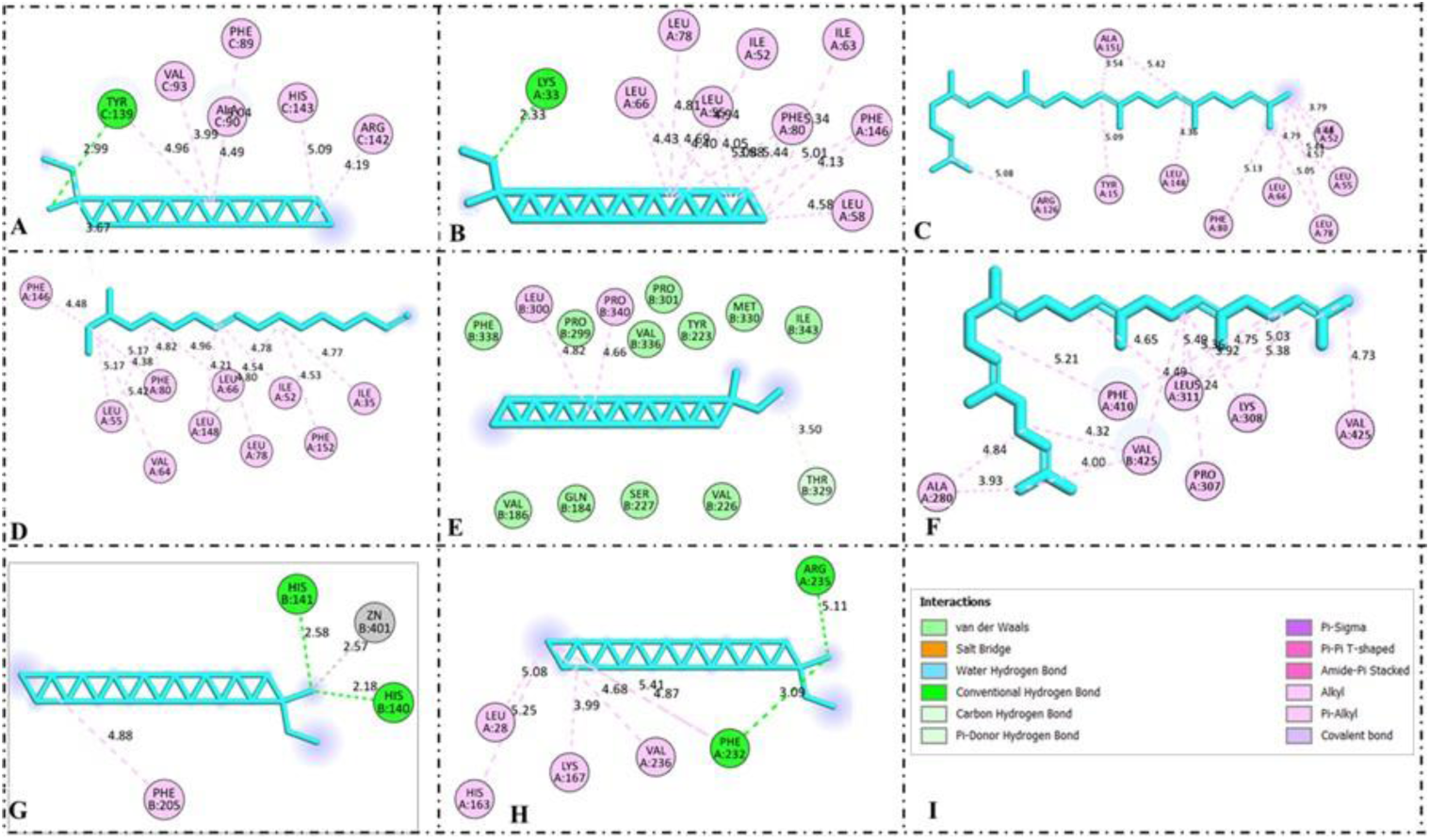
Two-dimension (2D) molecular docking results. Binding modes of compounds of ethyl acetate fraction onto binding-receptor pockets of the selected target protein. A) Methyl stearate bind BCL2, B), C) & D) Methyl stearate, squalene and (8Z)-14-methylhexadec-8-enal bind CDK2, E) Methyl stearate bind CDK1, F) Squalene bind P21, G) Methyl stearate bind HDAC1, and H) Methyl stearate bind CCND1

### 4.7 Validation of *M. edulis* fractions molecular targets through gene expression analysis by RT-qPCR

To determine the expression levels of 8 different genes that were selected from the top 30 hub genes and literature due to their established role in the regulation of cell cycle and apoptosis [21, 25, 77, 1, 86], the Livak method (ΔΔ*^CT^* method) was used to determine the relative gene expression with GAPDH gene being used as a house-keeping control gene to normalise the relative expressions. In HeLa cells treated with ethyl acetate fraction of *M. edulis*, among the 8 genes assessed, 3 genes were down-regulated while 4 genes were upregulated (Figure 17). Significantly downregulated genes are B cell lymphoma 2 (BCL2; p*<*0.0069), Cyclin-dependent kinase 2 (CDK2; p *<*0.0102), and Histone Deacetylase (HDAC1; p *<*0.0012) in comparison to negative control. On the other hand, significantly upregulated genes are Tumor suppressor (TP53; p *<*0.0003), Caspase-9 (CASP9; p *<*0.0001), CDK inhibitor (P21; p *<*0.0001), and Cyclin D1 (CCND1; p *<*0.0001) in comparison to the negative control. The gene expression level of cyclin-dependent kinase 1 (CDK1) was not significantly expressed (p *>*0.05).

**Fig. 17.**
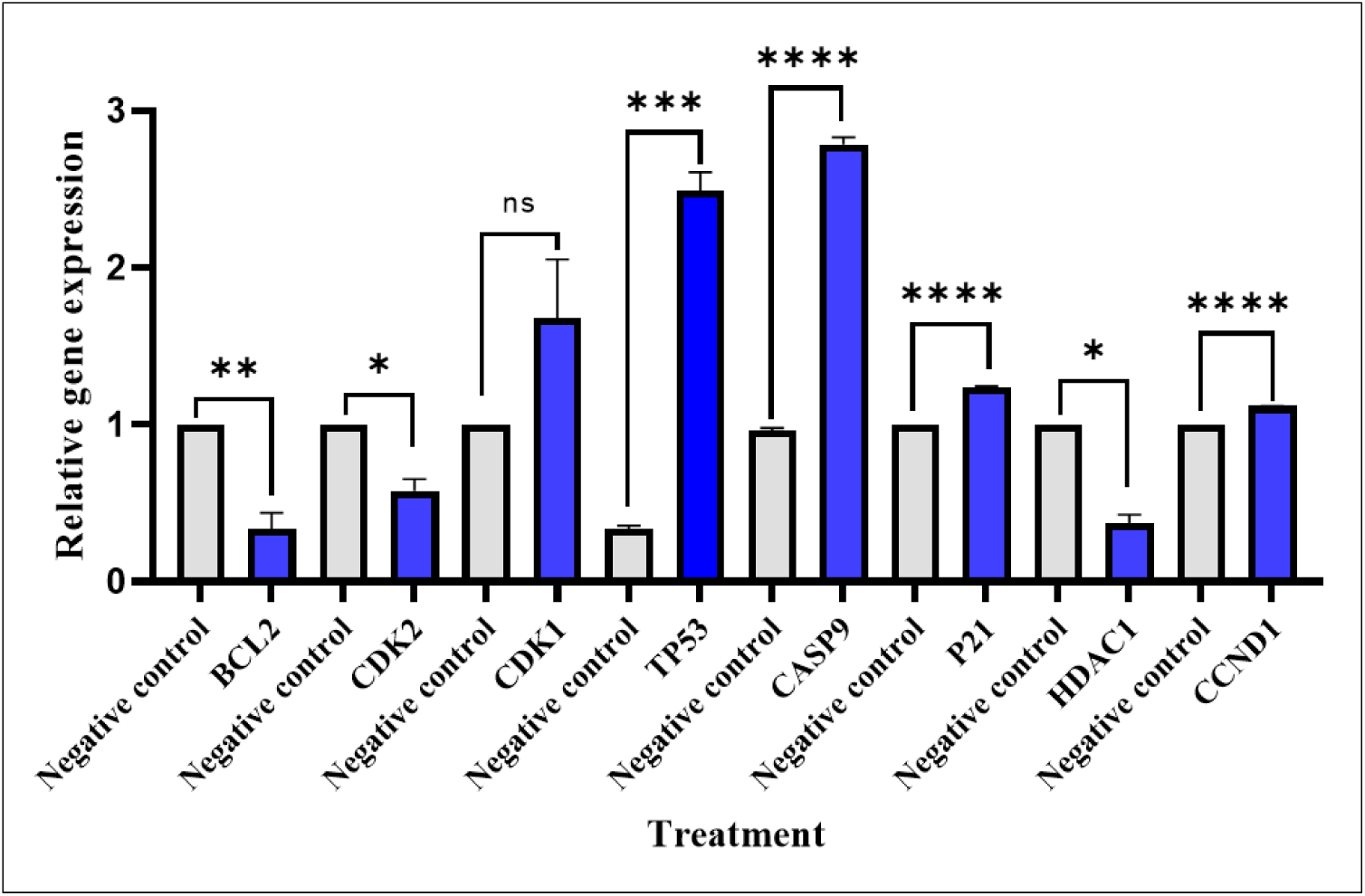
Effect of ethyl acetate fraction on expression levels of key genes in HeLa cells. Significance levels: ns, *p >* 0.05; *, *p <* 0.05; **, *p <* 0.01; ***, *p <* 0.001; ****, *p <* 0.0001.

In HeLa cells treated with hexane fraction of *M. edulis*, 4 genes were significantly downregulated, including BCL2; (p *<*0.0296), CDK2 (p *<*0.0001), HDAC1 (p *<*0.0018), and CCDN1 (p *<*0.001) in comparison to negative control while CDK1 was significantly upregulated with a p-value of 0.0001 as compared to negative (Figure 18). TP53 (p *>*0.2196), CASP9 (p *>*0.2112) and P21 (p *>*0.1458) genes were not significantly deregulated compared to negative control.

**Fig. 18.**
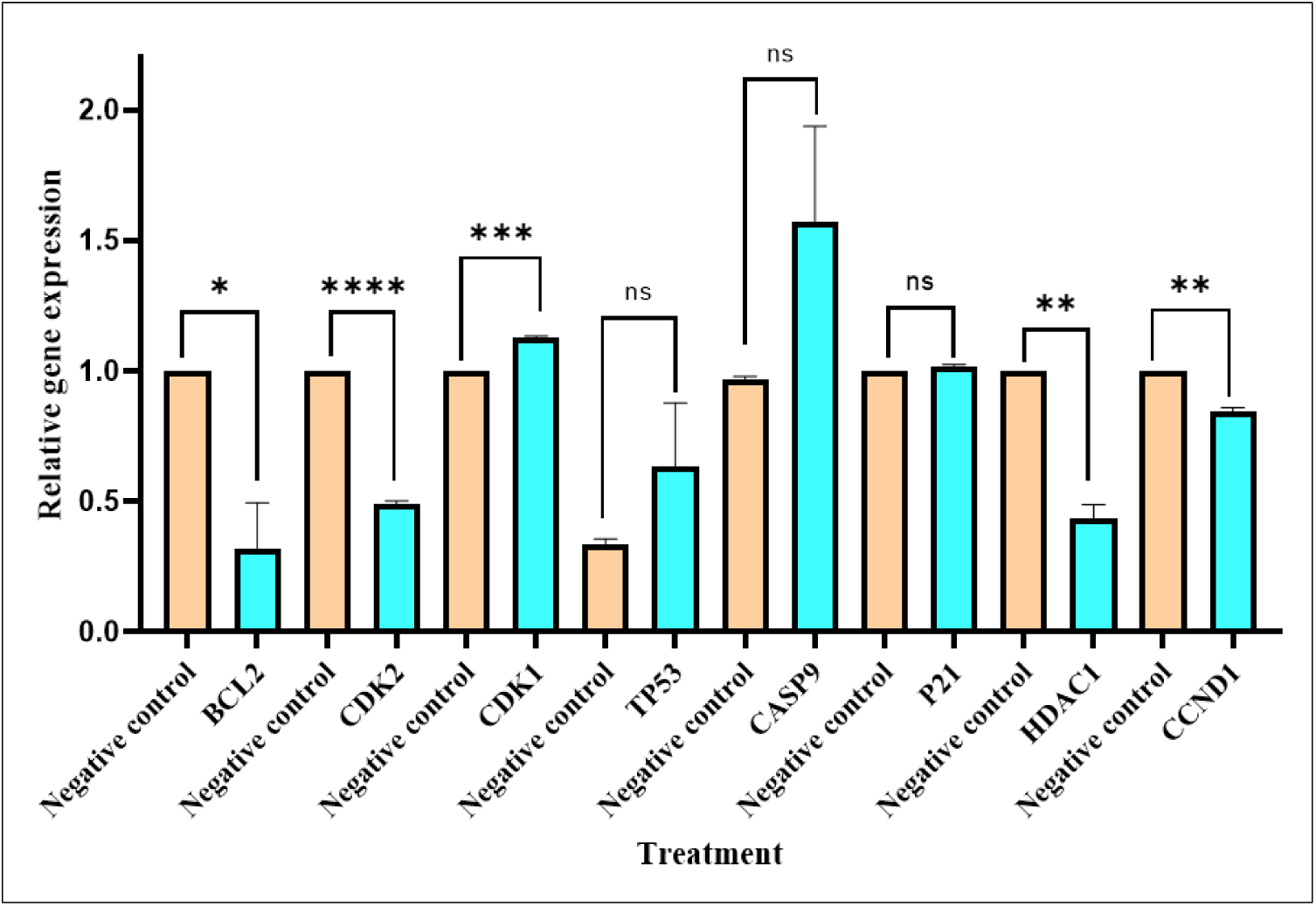
Effect of hexane fraction on expression levels of key genes in HeLa cells. Significance levels: ns, *p >* 0.05; *, *p <* 0.05; **, *p <* 0.01; ***, *p <* 0.001; ****, *p <* 0.0001.

## 5 Discussion

Natural products and their derivatives from medicinal plants are known for their robust biological activities such as antioxidants, antimicrobial, antidiabetic, antimalarial, anticancer properties [40, 53, 56], activity against cardiovascular diseases, Alzheimer’s disease, and other infections [13, 74]. *Maerua edulis* is a medicinal plant that has been reported to possess various therapeutic potentials, including anticancer activity against human leukemic Jurkat-T cell lines [70]. Ethnobotanically, *M. edulis* has been used to treat ailments such as eye infections, stomach aches, infertility in women, wounds, fungal infections, rheumatic swellings, cough, tuberculosis, and sexually transmitted diseases [70, 50]. However, scientific validation of its anticancer activity, particularly against cervical cancer, remains speculative. Therefore, this study chose to fill this lacuna by evaluating the antiproliferative and cytotoxic effects of *M. edulis*, including its putative mechanisms of action. Importantly, this is the first study to do so on the cervical cancer cell line model, to the best of our knowledge.

The initial screening of *M. edulis* extract and its solvent fractions against HeLa cells revealed that ethyl acetate and hexane fractions inhibited the growth and proliferation of HeLa cells remarkably, with hexane fraction being the most active followed by ethyl acetate fraction. The results further demonstrated that the cell viability was inhibited in dose-dependent manner for both fractions. The antiproliferative activity of the prioritized fractions on HeLa cells revealed that both fractions exhibited strong bioactivity. The guidelines of the US National Cancer Institute (NCI) classified IC_50_ value of a crude extract less than 30 *µg/mL* after 48 h of exposure to cancer cells as a potential cytotoxic candidate [38]. Therefore. Ethyl acetate fraction with IC_50_ above 30 *µg/mL* is considered to have moderate cytotoxic effects, while Hexane fraction exhibited an IC_50_ value estimated to be below 30 *µg/mL* and is thus considered highly cytotoxic.

We further evaluated *M. edulis* fractions for safety by testing them against viable noncancerous cells [5, 63]. Ethylacetate and hexane fractions of *M. edulis* that exhibited good antiproliferative activity against HeLa cells showed minimal cytotoxic effects on noncancerous Vero cells. Furthermore, selectivity index (SI) of *M. edulis* fractions was calculated to measure in vitro toxicity. Hypothetically, SI greater than 2 suggests the desired selectivity against cancer cells while preserving normal cells. The SI of both fractions evaluated after 48 h was greater than 5; meanwhile, doxorubicin had a SI of less than 2. These findings indicate that *M. edulis* fractions had a good in vitro anticancer activity, within acceptable in vitro toxicity threshold, thus giving a promise as a potential source for active yet safe anti-cervical cancer products. A higher SI value hypothetically predicts higher efficacies of extracts in an in vivo treatment model since normal cells will require a higher dose to die, while cancer cells will die at a lower dose [53].

The cytotoxic effects of *M. edulis* were phenotypically visualized on HeLa cells treated with *M. edulis* fractions compared to untreated cells as our negative control. It was observed that the cell morphology changed from a long-elongated shape to small round and/or spherical shape, shrunk shapes, and with blebbing which ultimately caused reduced cell viability. Similar results were reported when HeLa cells were exposed to *Hexogonia glabra* and *Agaricus bisporus* extracts, albeit at higher concentrations, but under the same conditions [21, 84].

Cell migration is involved in several physiological processes as a normal event. However, uncontrolled migration can result in cancer metastasis which accounts for 90% cause of human cancer-associated deaths [32]. Metastasis is a complex, multistep process that involves cell migration and invasion. Metastasis is frequently upregulated in 90% of cancer [91]. In the present study, ethyl acetate and hexane fractions of *M. edulis* significantly impaired the migratory capability of HeLa cells in an in vitro wound healing model. These findings are similar to studies reported by Kumar et al. (2022) on the ability of methanolic *Azadirachta indica* to inhibit the migrative ability of HeLa cells. These findings unveiled the potential of *M. edulis* extracts as a potential option for reducing the metastatic ability of cancer cells.

The preliminary qualitative and GC-MS phytochemical profiling of *M. edulis* revealed the presence of terpenoids, tannins, flavonoids, fatty acids and steroids as dominant phytochemicals. These phytochemicals have been reported to have arrays of medicinal properties, including antioxidant, anticancer, antimicrobial and antimalarial activities [18, 3, 58]. The presence of these phytochemicals in bioactive *M. edulis* extract fraction can partly or wholly explain the observed anticervical activity. Terpenoids and their derivatives (including squalene and stigmasta-3,5-diene) have anticancer activity, with several studies reporting terpenoids’ in vitro antiproliferative effects on pancreatic, gastric and colon carcinomas through cell cycle arrest, apoptosis induction and inhibition of metastasis [18, 32, 47, 36]. Tannins have antiapoptotic, antiaging, anticancer, antiinflammation and antiatherosclerosis activities [89, 24]. On the other hand, fatty acids (including (8Z)-14-Methyl-8-hexadecenal and methyl stearate) have been documented to inhibit or/and prevent cancer development [6, 76]. Flavonoids have strong anticancer activity by modulating multiple molecular processes, including induction of apoptosis, cell cycle arrest and autophagy, thus suppressing cancer cell proliferation and progression [4, 67]. Additionally, flavonoids have been reported to possess antitumor effects in high-grade adult-type diffuse glioma via regulating autophagy-related pathways [71]. Flavonoids inhibit Glioblastoma (GBM) and astrocytoma cell migration and invasion through inhibition of MMP-9, Erk and p38 protein activities [35]. Flavonoids also modulate the MAPK, p53 and NF-B pathways and their downstream proteins, such as the Bax/Bcl-2, caspase-3 and −9 in tumor cells [41]. Steroids and their derivatives have diverse biological activities, including anticancer, antibacterial, antifungal, antioxidant activities, among others [8, 19, 69, 51]. Taken together, the phytochemical profiling of *M. edulis* extract fractions showed that the medicinal plant is an excellent source of secondary metabolites.

Due to the presence of these bioactive phytochemicals, *M. edulis* can thus be considered an important source of therapeutic agents. However, it is suggested that further work on bioassay-guided isolation be carried out to isolate, purify and characterize the specific active phytochemicals of *M. edulis*.

Gene ontology (GO) enrichment and analysis depicted that *M. edulis* phytochemicals targeted key biological processes (BP), cellular component (CC) and Molecular Functions (MF), thus supporting and confirming their potential role in arresting the proliferation of cervical cancer model cell line. The response to chemical stimuli, organic substances, and oxygen-containing compounds are implicated in cell death and cell transformation into malignancies [34, 78, 66]. Regulation of programmed cell death and biological quality are normal biological processes in viable cells, which balance cell death with survival rate [17, 73]. They play vital roles in the definitive decision of tumor cell fate and most cancers have been linked to programmed cell death regulation [28]. Therefore, *M. edulis* by targeting these BP terms may indicate that it plays a significant role in the observed anti-cervical cancer effects in this study.

KEGG enrichment and analysis revealed that *M. edulis* may play an anti-cervical cancer role by regulating multiple biological pathways such as pathway in cancer, proteoglycans in cancer, PI3K-Akt1 signalling pathway, neuroactive ligand-receptor interaction, metabolic pathways, and retrograde endocannabinoid signalling which are implicated in the regulation of cell proliferation, cell division, metastasis, angiogenesis, differentiation, drug resistance, drug metabolism, cell population and apoptotic processes. These biological processes have negative and positive effects on the progression and regulation of tumorigenesis [29, 22]. Proteoglycans in cancer play crucial roles as a repertoire of molecular interactions implicated in tumor progression [16]. It has multifunctional roles in both normal and pathological processes such as morphogenesis, wound healing, inflammation and tumorigenesis [7, 82, 2]. A remodeled matrix in cancer stroma is linked with regulation of cancer cell phenotype and aggressiveness [82]. PI3K-Akt1 signalling pathway is crucial for numerous cellular functions, including cell proliferation, survival, adhesion, migration and metabolism [68, 20]. In cancer cells, PI3k-Akt1 signalling pathway regulates cancer cell proliferation, apoptosis, growth, transformation, and drug resistance [23, 37]. Studies have shown that the regulation of the PI3k-Akt1 signalling pathway inhibited cervical cancer growth, proliferation, apoptosis, invasion, and reversal of drug resistance by Chinese medicine monomers, SKA3 inhibitors and parthenolide [26, 45, 27]. Metabolic pathways and retrograde endocannabinoids play key roles in cell death and survival [10, 61]. These pathways are critical for tumor growth and survival by enhancing their biological needs such as need for proteins, lipids and nucleic acid [65, 42, 61]. Studies have shown that cervical cancer reprogrammed its metabolic pathways to meet its energy requirement; thus, modelling cancer’s metabolic pathways presents novel biomarkers for diagnosis and therapy [61, 42].

In this current study, we analyzed the phytochemical components and key molecular targets of *M. edulis* with the aid of GC-MS, protein-protein interactions and KEGG analyses. Based on this approach, we were able to identify 20 and 14 compounds in ethyl acetate and hexane extract fractions (among them; diisooctyl phthalate, squalene, stigmasta-3,5-diene, (8Z)-14-methylhexadec-8-enal, methyl stearate, and methyl-5,11,14,17-eicosatetraenoate and 8 molecular targets (BCL2, CDK2, CDK1, TP53, CASP9, P21, HDAC1, and CCDN1) that were further prioritized for molecular docking (MD) analysis. The docking results showed binding energies ranging between −8.5 and −6.0 kcal/mol, suggesting good binding activity between these potentially active compounds and their potential molecular targets.The high binding energies can be attributed to favourable interactions between these phytochemical molecules and their molecular targets, which included conventional hydrogen bonds, Van der Waals forces, pi-donor hydrogen bonds, pi-Alkyl interactions, pi-sigma and carbon-hydrogen bonds and other hydrophobic interactions. The results of MD suggest that diisooctyl phthalate, squalene, stigmasta-3,5-diene, (8Z)-14-methylhexadec-8-enal, methyl stearate, and methyl-5,11,14,17-eicosatetraenoate may serve as potential key active compounds for the treatment of cervical cancer. We further validated the docking results by redocking with the native ligands of the target proteins, and it was noted that the binding energies ranged between −8.7 to −4.2 kcal/mol. These results suggest that *M. edulis* phytochemicals are promising inhibitors for the stated cervical cancer molecular targets.

To further affirm and validate targets that could potentially have been deregulated by extract treatment and thus accounting for the observed phenotype, we tested select candidate targets for gene expression by RT-qPCR approach. We observed downregulation of B cell lymphoma 2 (BCL2), Cyclin-dependent kinase 2 (CDK2), and Histone Deacetylase (HDAC1) genes after treatment of HeLa cells with *M. edulis* extract fractions. In cancer development, these genes are generally upregulated. BCL2 is a key regulator of apoptosis [39], and its upregulation is associated with the development of cervical cancer by inhibiting apoptosis [21]. CDK2 is crucial for cell proliferation via regulation of DNA synthesis [88]. In cervical cancer, CDK2 is overexpressed to inhibit cell cycle arrest [88] hence, promoting cancer cell proliferation. Inhibiting CDK2 expression by *M. edulis* is a promising mechanistic intervention against cervical cancer. HDAC1 expression promotes proliferation and invasion of osteosarcoma cells via inhibiting the Micro-RNA-326 transcription gene [83]. The observed down-regulation of HDAC1 upon treatment of HeLa cells by *. edulis* gives a mechanistic explanation of the anticervical cancer activity of *M. edulis* extract fractions. Furthermore, *M. edulis* extract induced an upregulation of Tumor (suppressor) protein p53 (TP53), Caspase-9 (CASP9), CDK inhibitor (P21), and Cyclin D1 (CCND1) genes. TP53 is a key pro-apoptotic factor and tumour inhibitor, and thus many antitumor drugs exerts their anticancer activity by targeting TP53-related signalling pathways [86]. In cancer cells, TP53 expression is usually suppressed, thus causing loss of the ability to destroy tumor cells through apoptosis, and this suppression is found in 60% of human cancers [73]. P21 is an inhibitor of CDK complexes and its upregulation is mediated by TP53-dependent and TP53-independent mechanisms and thus it plays a key role in G1 and G2 cell cycle arrest in response to DNA damage [64, 79]. In this study, *M. edulis* extract fraction treatment induced a significant upregulation of CDK2 and CDK1 inhibitor (P21), which can partly or wholly offer mechanistic explanation of the observed growth inhibition of treated HeLa cells. CCND1 is a crucial mediator of cell cycle progression [86]; its expression accelerates cell proliferation via promoting cell cycle progression. Therefore, CCND1 is highly overexpressed in multiple cancers, including cervical cancer [52]. Its downregulation causes abnormal cell proliferation and promotes the development of cancer [81]. Caspase 9 is considered a key enzyme that triggers a cascade of events that result in apoptosis [15]. *M. edulis* promotes overexpression of Caspase 9, and this may offer an additional mechanistic explanation of the significant in-vitro antiproliferative effects on HeLa cells after 48 hr of extract treatment.

## 6 Conclusion

*Maerua edulis* contains a diverse array of phytochemicals that can be associated with the observed robust antiproliferative, anti-migratory activities and deregulation of critical cancer driver and suppressor target proteins. In silico studies showed that *M. edulis* 6 predominant phytochemicals (diisooctyl phthalate, squalene, stigmasta-3,5-diene, (8Z)-14-methylhexadec-8-enal, methyl stearate, and methyl-5,11,14,17-eicosatetraenoate) demonstrated putative multi-faceted molecular mechanistic actions by modulating multiple biological functions and signalling pathways associated with tumorigenesis. Furthermore, these phytochemicals have a high binding affinity to the receptor-binding pocket of apoptotic- and cell-cycle-related genes and we could recapitulate these in silico findings by demonstrating phenotypic gene expression deregulation upon treating HeLa cells with *M. edulis* extracts. Taken together, these phenotypic and mechanistic in silico and in vitro findings confirm the potential of *M. edulis* extract as potential alternative source for cervical cancer therapy. However, further compound isolation, in vivo, and protein studies are recommended to assess its efficacies and safety.

## Supporting information

Suplementary Sheet 1

## Supplementary Materials

Supporting information and data can be retrieved from the journal’s supplementary materials section.

## Author Contributions

I.J.L.L.: Conceptualisation, Methodology, Investigation, Data curation, Writing—Original draft preparation. R.L.: Methodology, Reviewing, Proofreading.F.M.M.: Methodology, Reviewing, Proofreading. S.W.K.: Methodology, Reviewing, Proofreading. J.M.M, F.M.M & V.K: Reviewing and Proofreading. D.W.N.: Conceptualisation, Supervision, Reviewing, Editing. S.N.N.: Conceptualisation, Sample acquisition, Methodology, Supervision, Reviewing, Editing. All authors have read and approved the final manuscript.

## Institutional Review Board Statement

The go-ahead for the study was granted by the Kenya Medical Research Institute (KEMRI) Review Board; Scientific Ethical Review Unit (SERU)(REF: KEMRI/SERU/CTMDR/104/4466).

## Informed Consent Statement

Not applicable.

## Data Availability Statement

The data presented in this study are available upon request from the corresponding authors.

## Acknowledgements

We acknowledge the African Union (AU) through Pan African University for Basic Sciences, Technology and Innovation (PAUSTI), the Centre for Traditional Medicine and Drug Research, Kenya Medical Research Institute (KEMRI), and Jomo Kenyatta University of Agriculture and Technology (JKUAT) for providing laboratories and resources. Special thanks to Mercy Jepkorir, Wesley Kanda, and Amel Elbasyouni for their assistance during lab work.

## Footnotes

1 https://pubchem.ncbi.nlm.nih.gov/

2 http://www.swissadme.ch/index.php

3 https://biosig.lab.uq.edu.au/pkcsm/prediction

4 http://www.swisstargetprediction.ch/database

5 https://bindingdb.org

6 https://sea.bkslab.org/

7 https://www.uniprot.org

8 https://www.genecards.org/

9 https://www.disgenet.org/

10 https://www.omim.org/

11 https://pharos.nih.gov/diseases/

12 https://bioinformatics.psb.ugent.be/webtools/ven

13 https://cn.string-db.org

14 https://ge-lab.org/go/

15 https://www.rcsb.org/

