## Supplementary material for "Antiproliferative Activity, Phytochemistry, Network Pharmacology, Molecular Docking and Gene Expression Analysis of *Maerua edulis* Extracts against Human Cervical Cancer Cell Line": Suplementary Sheet 1

Supplementary Table 1: Forward primers and reverse primers of genes used for RT-qPCR analysis.

| Gene | Forward primer (5’-3’) | Reverse primer (5’-3’) |
| --- | --- | --- |
| BCL2 | GGCCTCAGGGAACAGAATGAT | TCCTGTTGCTTTCGTTTCTTTC |
| CDK1 | CGCGGATCCATGGAAGATTATACCAAAA | CCGGAATTCCTACATTCTTAATCTGAT |
| CDK2 | TCTTTGCTGAGATGGTGACTCG | TGTTAGGGTCGTAGTGCAGC |
| HDAC1 | TACGACGGGGATGTTGGAAA | ATTGGCTTTGTGAGGGCGAT |
| TP53 | CTTCGAGATGTTCCGAGAGC | CTTCGAGATGTTCCGAGAGC |
| CASP9 | CCTGCCCGCTGTTTGGA | GCTGGGAAATGGGGAGACAA |
| P21 | GCGACTGTGAGCTAATG | TTAGAAGCTTGGCAAAGGGC |
| CCND1 | GAGGCGGAGGAGAACAAACA | GGAGGGCGGATTGGAAATGA |
| GAPDH | AGACAGCCGCATCTCTTG | TGACTGTGCCGTTGAACTTG |

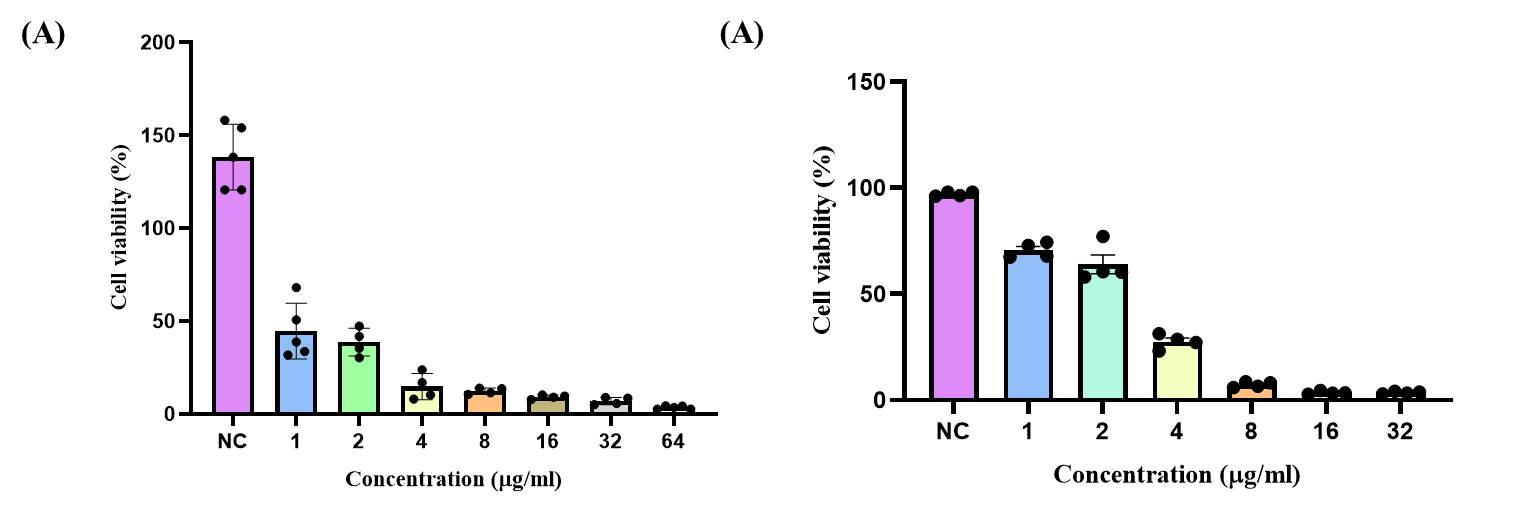

Supplementary Figure 1: In-vitro antiproliferative and cytotoxic effects of Doxorubicin extracts on HeLa and Vero cells. (A) dose-dependent effects of Doxorubicin on HeLa. (B) dose-dependent effects of Doxorubicin on Vero cells after 48 h of treatment. NC-negative control.

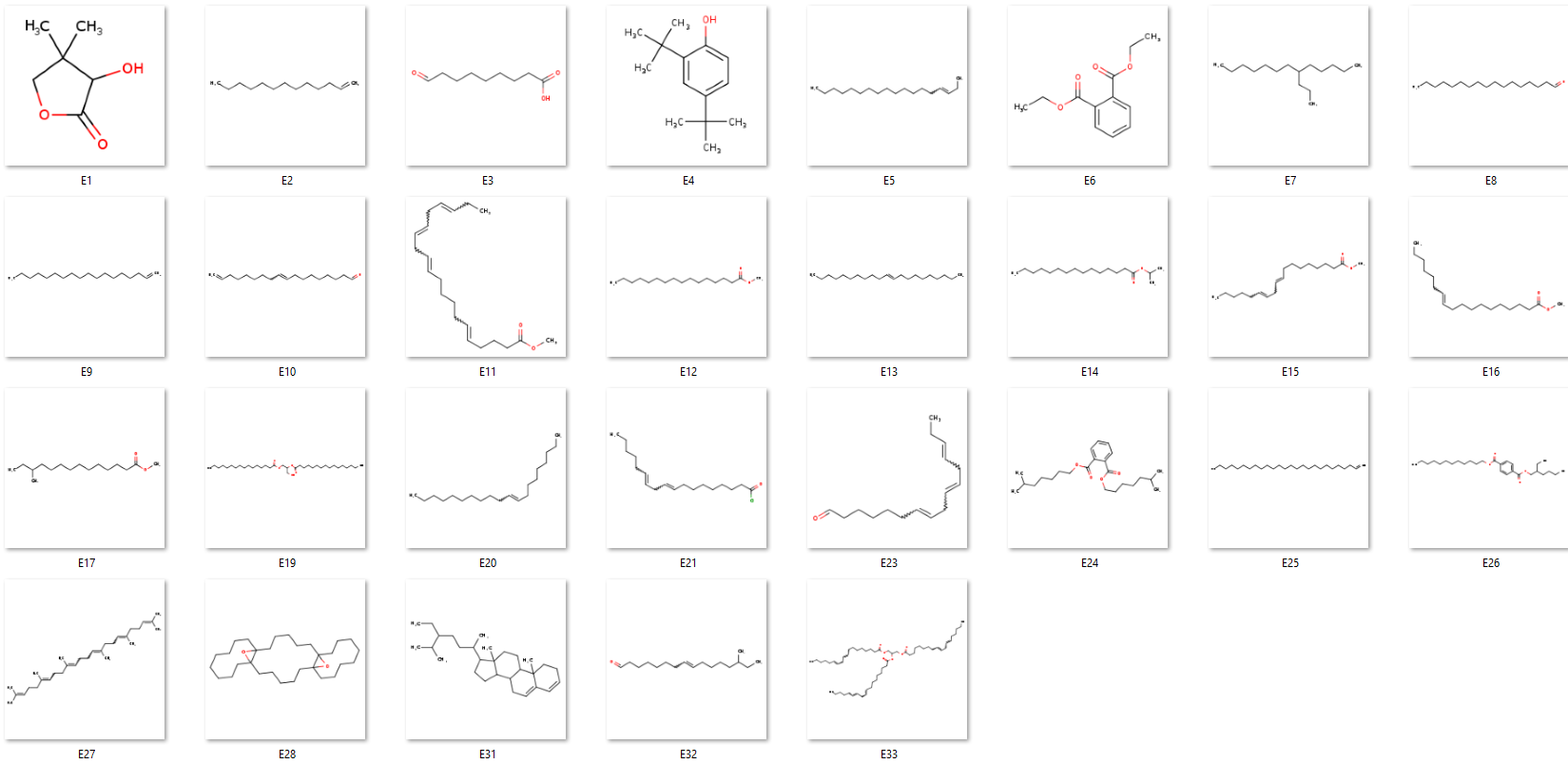

Supplementary Figure 2: The letter E1-E33 presents the 2D structure of the phytochemicals identified in ethyl acetate fraction of *M. edulis* (For Details names, refers to the manuscript Table 2).

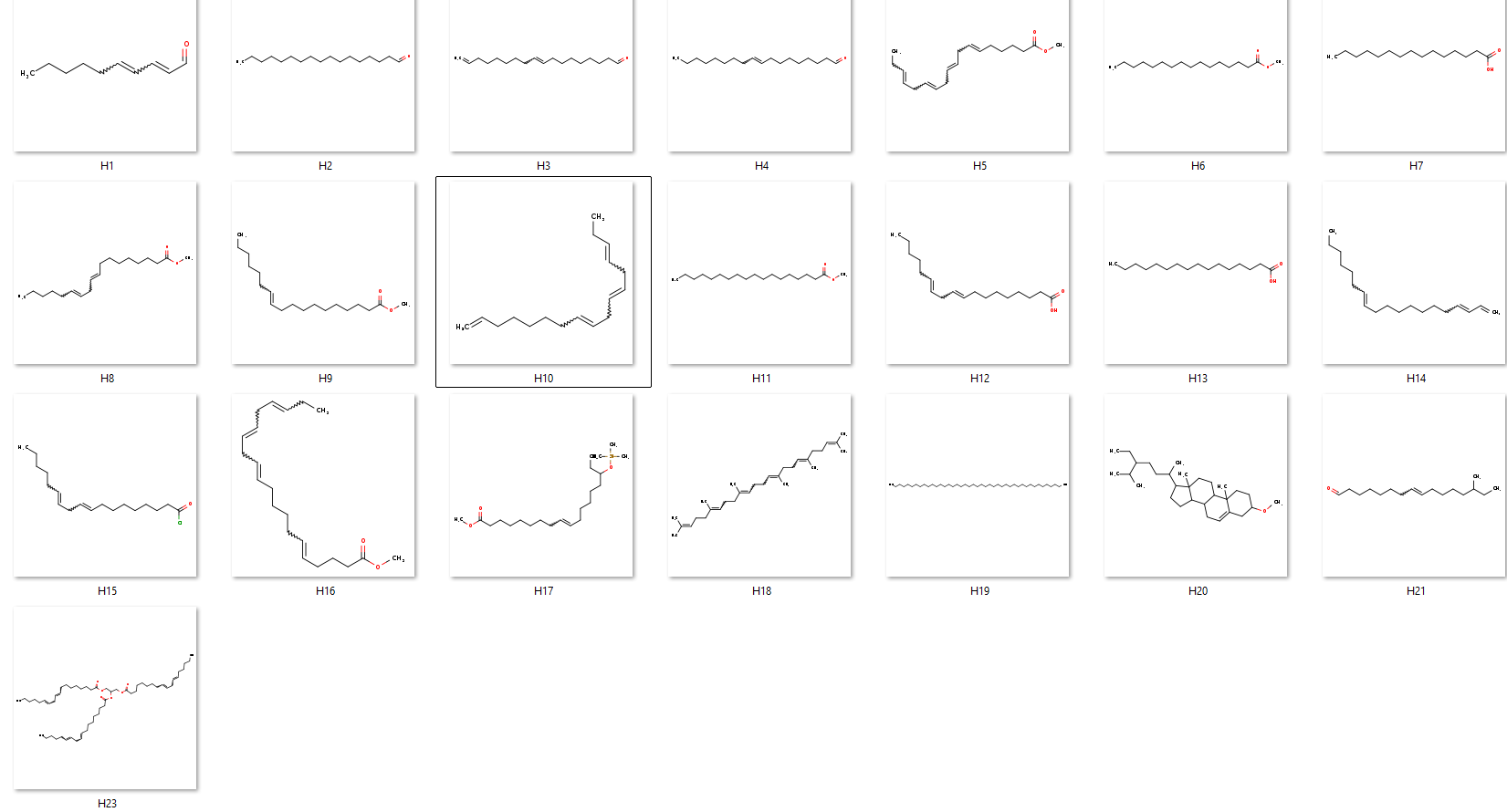
Supplementary Figure 3: The letter H1-H23 presents the 2D structure of the phytochemicals identified in hexane fraction of M. edulis (For Details names, refers to the manuscript Table 3).

Supplementary Table 2: Prediction of physiochemical properties for M. edulis ethyl acetate fraction compound based on Lipinski (majorly considered here), Veber, Egan, Muegge, and Ghose rules.

| Ligands  Number | Physical and chemical properties | | | | | | Lipinski  Violations | BBB |
| --- | --- | --- | --- | --- | --- | --- | --- | --- |
|  | MW (g/mol) | Molar refractive index | Rotatable  bonds number | LogP (Octanol/Water) | H-bond  Acceptors Number | H- bonds donors Number | Categorical  (Yes/No) | |
| Threshold | ≤500 | 40≤MR≤130 | ≤10 | ≤5 | ≤10 | ≤5 | Yes/No | Yes/No |
| E1 | 130.14 | 31.03 | 0 | 0 | 3 | 1 | Yes | Yes |
| E2 | 182.35 | 64.13 | 10 | 5.52 | 0 | 0 | Yes | Yes |
| E3 | 172.22 | 47.35 | 8 | 1.29 | 3 | 1 | Yes | Yes |
| E4 | 206.32 | 67.01 | 2 | 3.87 | 1 | 1 | Yes | Yes |
| E5 | 252.48 | 88.17 | 14 | 6.77 | 0 | 0 | **Yes** | **No** |
| E6 | 222.24 | 58.61 | 6 | 2.39 | 4 | 0 | Yes | Yes |
| E7 | 226.44 | 79.03 | 12 | 6.44 | 0 | 0 | Yes | **No** |
| E8 | 254.45 | 84.03 | 15 | 4.55 | 1 | 0 | Yes | **No** |
| E9 | 252.48 | 88.17 | 15 | 6.77 | 0 | 0 | Yes | **No** |
| E10 | 264.45 | 87.89 | 15 | 4.59 | 1 | 0 | Yes | **No** |
| E11 | 318.49 | 102.45 | 15 | 4.98 | 2 | 0 | **Yes** | **No** |
| E12 | 270.45 | 85.12 | 15 | 4.44 | 2 | 0 | Yes | **No** |
| E13 | 294.56 | 102.59 | 17 | 7.46 | 0 | 0 | Yes | **No** |
| E14 | 284.48 | 89.92 | 15 | 4.67 | 2 | 0 | Yes | **No** |
| E15 | 294.47 | 93.78 | 15 | 4.7 | 2 | 0 | Yes | **No** |
| E16 | 296.49 | 94.26 | 16 | 4.8 | 2 | 0 | **Yes** | **No** |
| E17 | 256.42 | 80.31 | 13 | 4.19 | 2 | 0 | Yes | **No** |
| E19 | 568.91 | 174.09 | 34 | 6.27 | 5 | 1 | **No** | **No** |
| E20 | 280.53 | 97.78 | 16 | 7.23 | 0 | 0 | Yes | **No** |
| E21 | 298.89 | 92.69 | 14 | 4.82 | 1 | 0 | Yes | **No** |
| E23 | 234.38 | 77.8 | 11 | 4.01 | 1 | 0 | Yes | **No** |
| E24 | 390.56 | 116.3 | 16 | 5.24 | 4 | 0 | Yes | **No** |
| E25 | 364.69 | 126.62 | 23 | 8.51 | 0 | 0 | **Yes** | **No** |
| E26 | 446.66 | 135.53 | 21 | 6.05 | 4 | 0 | Yes | **No** |
| E27 | 410.72 | 143.48 | 15 | 7.93 | 0 | 0 | Yes | **No** |
| E28 | 444.73 | 138.08 | 0 | 6.14 | 2 | 0 | Yes | **Yes** |
| E31 | 396.69 | 131.59 | 6 | 7.7 | 0 | 0 | Yes | **Yes** |
| E32 | 252.44 | 83.56 | 13 | 4.44 | 1 | 0 | **Yes** | **No** |
| E33 | 879.38 | 277.12 | 50 | 9.25 | 6 | 0 | **No** | **No** |

Supplementary Table 3: Prediction of physiochemical properties for M. edulis hexane fraction compound based on Lipinski (majorly considered here), Veber, Egan, Muegge, and Ghose rules.

| Ligands  Number | Physical and chemical properties | | | | | | Lipinski  Violations | BBB |
| --- | --- | --- | --- | --- | --- | --- | --- | --- |
|  | MW (g/mol) | Molar refractive index | Rotatable  bonds number | LogP (Octanol/Water) | H-bond  Acceptors Number | H- bonds donors Number | Categorical  (Yes/No) | |
| Threshold | ≤500 | 40≤MR≤130 | ≤10 | ≤5 | ≤10 | ≤5 | Yes/No | Yes/No |
| H1 | 152.23 | 49.44 | 6 | 2.878 | 1 | 0 | Yes | Yes |
| H2 | 254.45 | 84.03 | 15 | 6.0567 | 1 | 0 | Yes | No |
| H3 | 264.45 | 87.89 | 15 | 5.9988 | 1 | 0 | Yes | No |
| H4 | 266.46 | 88.37 | 15 | 6.2228 | 1 | 0 | Yes | No |
| H5 | 290.44 | 92.84 | 13 | 5.5249 | 2 | 0 | Yes | Yes |
| H6 | 270.45 | 85.12 | 15 | 5.6407 | 2 | 0 | Yes | Yes |
| H7 | 242.4 | 75.99 | 13 | 5.1622 | 2 | 1 | Yes | Yes |
| H8 | 294.47 | 93.78 | 15 | 5.9729 | 2 | 0 | Yes | No |
| H9 | 296.49 | 94.26 | 16 | 6.1969 | 2 | 0 | Yes | No |
| H10 | 232.4 | 81.94 | 11 | 5.9817 | 0 | 0 | Yes | No |
| H11 | 298.5 | 94.73 | 17 | 6.4209 | 2 | 0 | Yes | No |
| H12 | 280.45 | 89.46 | 14 | 5.8845 | 2 | 1 | Yes | Yes |
| H13 | 256.42 | 80.8 | 14 | 5.5523 | 2 | 1 | Yes | Yes |
| H14 | 262.47 | 92.03 | 14 | 6.9859 | 0 | 0 | Yes | No |
| H15 | 298.89 | 92.69 | 14 | 6.5653 | 1 | 0 | Yes | No |
| H16 | 318.49 | 102.45 | 15 | 6.3051 | 2 | 0 | Yes | No |
| H17 | 384.67 | 117.63 | 18 | 7.0269 | 3 | 0 | Yes | No |
| H18 | 410.72 | 143.48 | 15 | 10.605 | 0 | 0 | Yes | No |
| H19 | 605.16 | 208.82 | 40 | 17.0203 | 0 | 0 | No | No |
| H20 | 428.73 | 137.96 | 7 | 8.6789 | 1 | 0 | Yes | No |
| H21 | 252.44 | 83.56 | 13 | 5.6886 | 1 | 0 | Yes | No |
| H23 | 879.38 | 277.12 | 50 | 17.4251 | 6 | 0 | No | No |

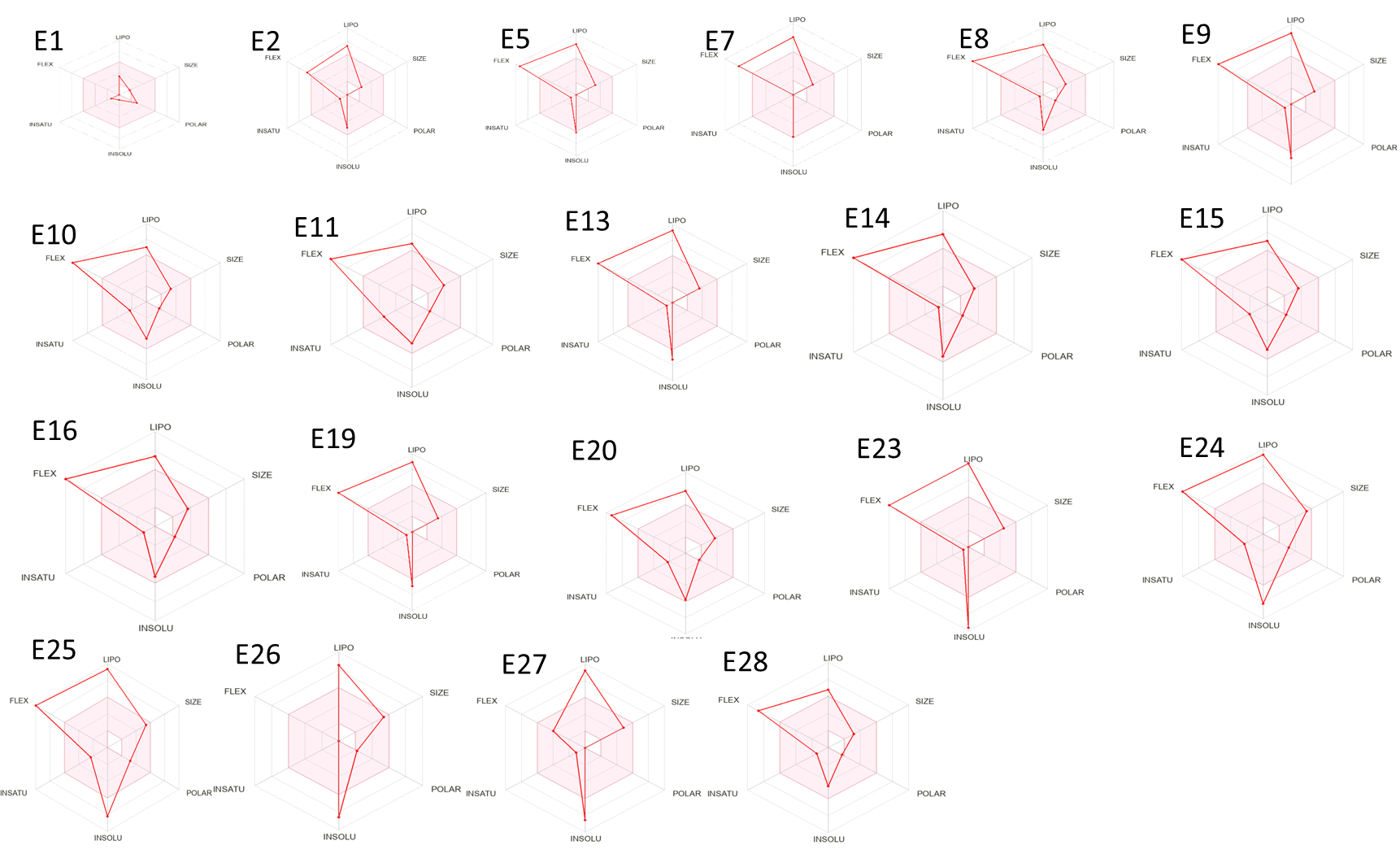

Supplementary Figure 4: Bioavailability radars of 14 *M. edulis* ethyl acetate fraction drug-like compounds, based on the six ideal physicochemical properties for oral bioavailability, namely, polarity (POLAR), lipophilicity (LIPO), saturation (INSATU), size (SIZE), flexibility (FLEX), and solubility (INSOLU) (Details name of E1-E28 refers to Manuscript Table 4.

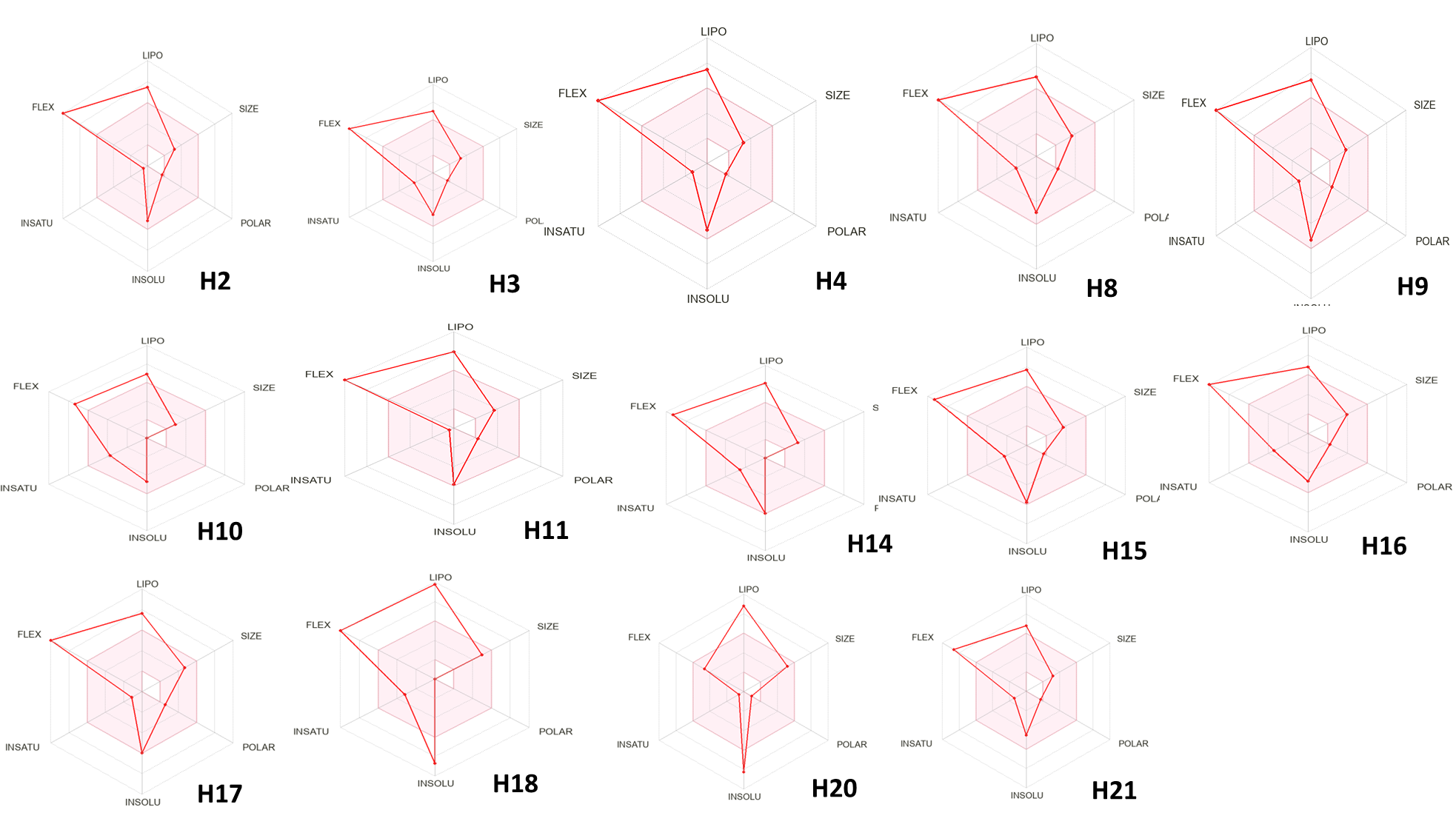
Supplementary Figure 5: Bioavailability radars of 14 *M. edulis* hexane fraction drug-like compounds, based on the six ideal physicochemical properties for oral bioavailability, namely, polarity (POLAR), lipophilicity (LIPO), saturation (INSATU), size (SIZE), flexibility (FLEX), and solubility (INSOLU), (Details name of H2-H21 refers to Manuscript Table 5).

Supplementary Figure 6. The shared common targets between M. edulis ethyl acetate and hexane fractions.

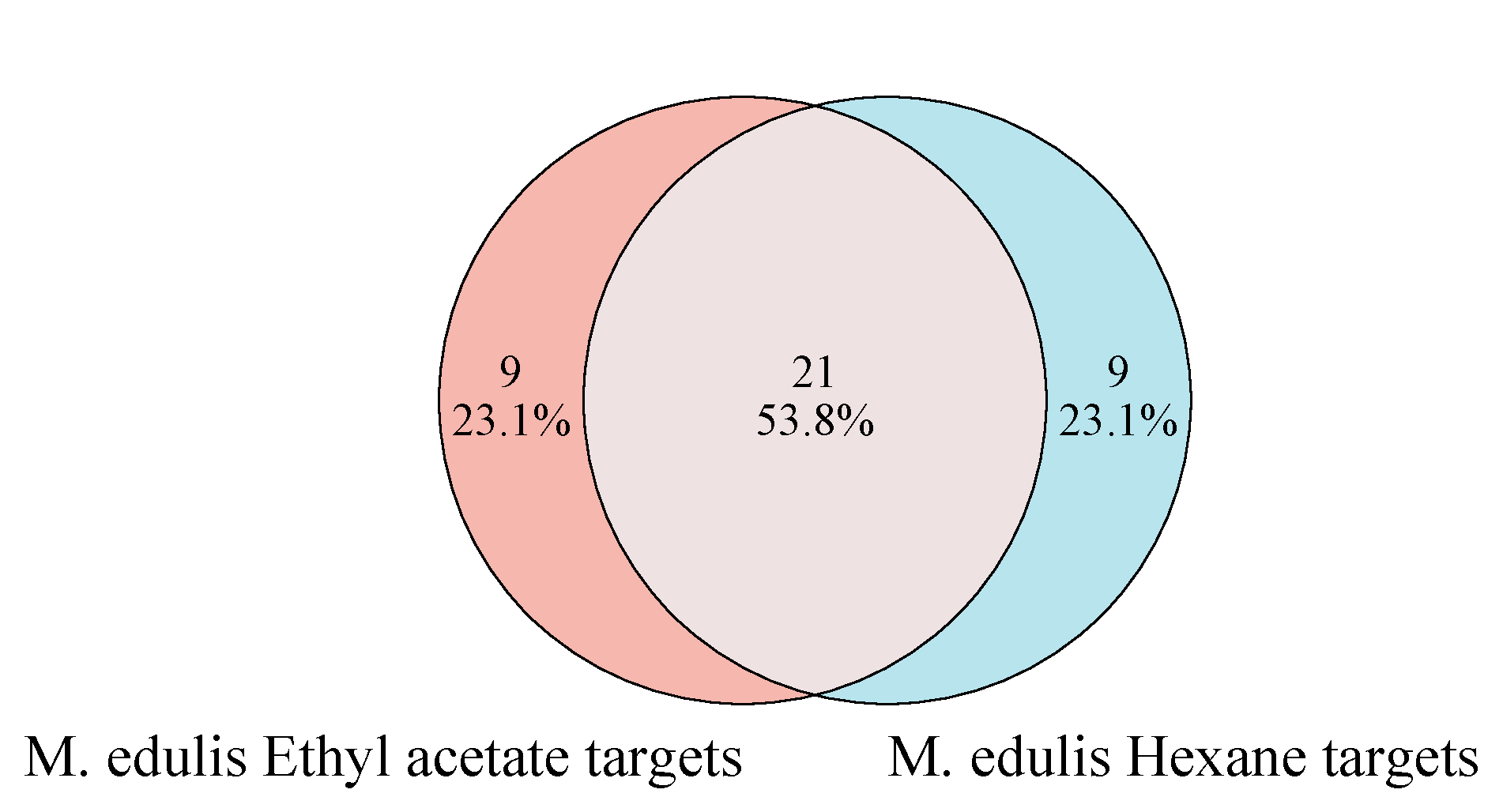

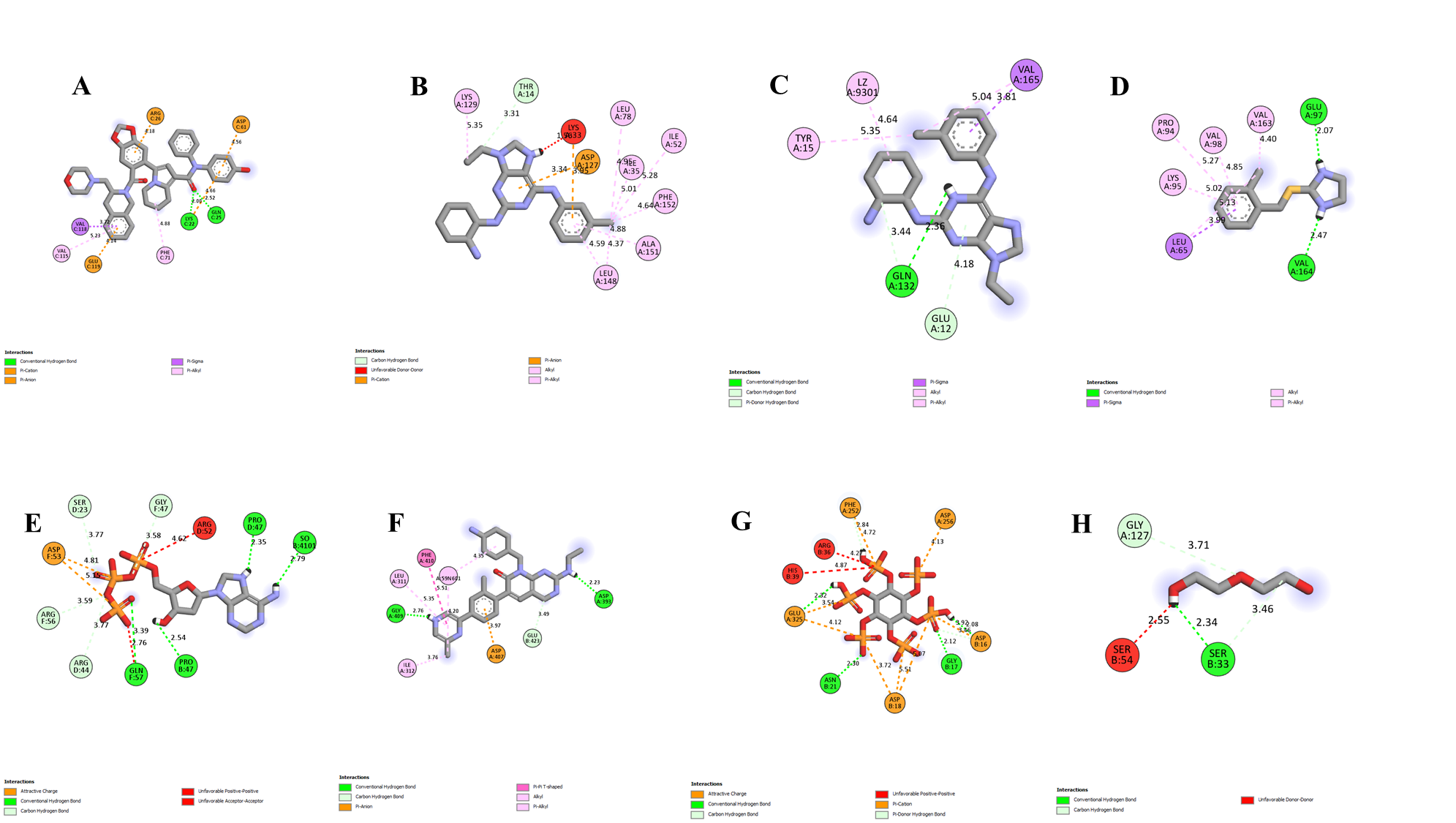

Supplementary Figure 7: Two-dimension (2D) molecular docking results. Binding modes of redocked native ligands onto binding-receptor pockets of the selected target protein. (A-BCL1; B-CDK2; C-CDK1; D-TP53; F-P21; G-HDAC1; H-CCND1).

Supplementary Table 4: The binding affinity in kcal/mol of the redocked native ligands with selected target proteins

| Protein/redocked ligand | BCL2 | CDK2 | CDK1 | TP53 | CASP9 | P21 | HDAC1 | CCND1 |
| --- | --- | --- | --- | --- | --- | --- | --- | --- |
| NL1 | **-8.7** |  |  |  |  |  |  |  |
| NL2 |  | **-8.7** |  |  |  |  |  |  |
| NL3 |  |  | **-7.6** |  |  |  |  |  |
| NL4 |  |  |  | **-5.2** |  |  |  |  |
| NL5 |  |  |  |  | **-7.6** |  |  |  |
| NL6 |  |  |  |  |  | **-8.5** |  |  |
| NL7 |  |  |  |  |  |  | **-5.5** |  |
| NL8 |  |  |  |  |  |  |  | -4.2 |
